# Temporal stimulus segmentation by reinforcement learning in populations of spiking neurons

**DOI:** 10.1101/2020.12.22.424037

**Authors:** Luisa Le Donne, Lik Chun Chan, Robert Urbanczik, Walter Senn, Giancarlo La Camera

**Author notes:** Space Biomedical Centre, University of Rome Tor Vergata, Rome, Italy (Luisa Le Donne). Deceased (Robert Urbanczik).

## Abstract

Learning to detect, identify or select stimuli is an essential requirement of many behavioral tasks. In real life situations, relevant and non-relevant stimuli are often embedded in a continuous sensory stream, presumably represented by different segments of neural activity. Here, we introduce a neural circuit model that can learn to identify action-relevant stimuli embedded in a spatio-temporal stream of spike trains, while learning to ignore stimuli that are not behaviorally relevant. The model uses a biologically plausible plasticity rule and learns from the reinforcement of correct decisions taken at the right time. Learning is fully online; it is successful for a wide spectrum of stimulus-encoding strategies; it scales well with population size; and can segment cortical spike patterns recorded from behaving animals. Altogether, these results provide a biologically plausible framework of reinforcement learning in the absence of prior information on the identity, relevance and timing of input stimuli.

## I. INTRODUCTION

Human beings have an unmatched ability to learn to abstract information from the environment, use this information to predict the consequences of their actions and develop appropriate behavioral strategies. A major component of this skill is the ability to identify the relevant cues and stimuli from the environment and determine what to do in response to them. Identifying relevant stimuli is a crucial aspect of many cognitive tasks including decision making. Apart from a few exceptions that, however, do not yet admit a biological implementation [1], the great majority of learning and decision making models [2, 3] typically assume that the relevant stimuli are known to the learning agent. In these models, learning refers to the process of associating known relevant stimuli with their predicted outcome for the purpose of making decisions (see e.g. [4] for a general framework). The underlying assumption is that relevant stimuli can be identified by a form of bottom-up, unsupervised preprocessing; for example, relevant stimuli are so because statistically different from other stimuli. Although this could be the case for perceptually salient stimuli that ‘stand out’ from the background, in many real-life situations it is not clear which segments in a continuous sensory stream are action relevant, and relevant segments may blend seamlessly into irrelevant ones. Here, we present a spiking network model (the learning agent, or ‘agent’ for short) able to detect relevant stimuli from the environment by trying to maximize the reward obtained for correct decisions at the right time. The agent is required to solve a detection and a decision task simultaneously: detect which stimuli are behaviorally relevant and decide what to do in response to them. The stimuli are modeled as spatiotemporal patterns of spike trains, and the agent has initially no knowledge of their identity, timing, or required response. Everything must be discovered by trial and error by exploiting reward feedback received after each decision. This is all the more challenging as relevant and non-relevant stimuli were built so as to have the same overall spike train statistics, so that different segments appear seamlessly embedded in a homogeneous input stream (see Figs. 2 and 4 for examples).

We emphasize that decisions are self-paced and never enforced in this paradigm, and no feedback is given in the absence of decisions. This is a particularly demanding requirement because the quiescent state (i.e., the network state where no decisions are ever taken) is a local maximum of the average reward, and can result in the pernicious behavior of forgoing decisions altogether. Here we show that, in a large variety of situations, our agent can learn to identify the relevant stimuli embedded in patterns of spike trains and take the correct decision in response to them. Although more slowly, the agent also learns to ignore non-relevant stimuli. We also show that the agent is equally successful at segmenting patterns of cortical spike trains recorded from behaving rodents. These results were achieved through a fully online synaptic plasticity rule modulated by reward and feedback from the population decision. Overall, our model provides a biologically plausible framework for learning to segment an input stream for the purpose of solving behavioral tasks, and may be one way to form concepts out of seemingly unstructured stimuli [5].

## II. RESULTS

### A. Segmentation task with multiple action choices

We trained an ensemble of populations of spiking neurons (the ‘agent’) to learn to detect specific stimuli embedded as segments of a continuous input stream. The agent had to produce a desired response to each of the detected segments. Each segment was a spatio-temporal pattern of spike trains lasting 500 ms (Fig. 1**a**). The segments were noisy ‘realizations’ of a handful of prototype stimuli (Fig. 1**b**), temporally joined together in random order to form a long, continuous stream of input patterns. In the default setting, we built each prototype by generating 50 independent Poisson spike trains, each with constant firing rate randomly chosen according to a uniform distribution between 2 and 24 spikes/s. Each prototype stimulus was randomly assigned to being either relevant or non-relevant for the task, a property inherited by their noisy realizations. The segments were obtained by jittering the spike times of the prototypes by a random amount of the order of 10 ms (see Fig. 1**b** and Sec. IV D of Methods for details), but we will show later that the model is robust to much longer jitter times (see Fig. 1**c** for an example segment with 100 ms jitter). In the following, we shall use the term ‘segment’ or ‘stimulus’ interchangeably.

**FIG. 1.**
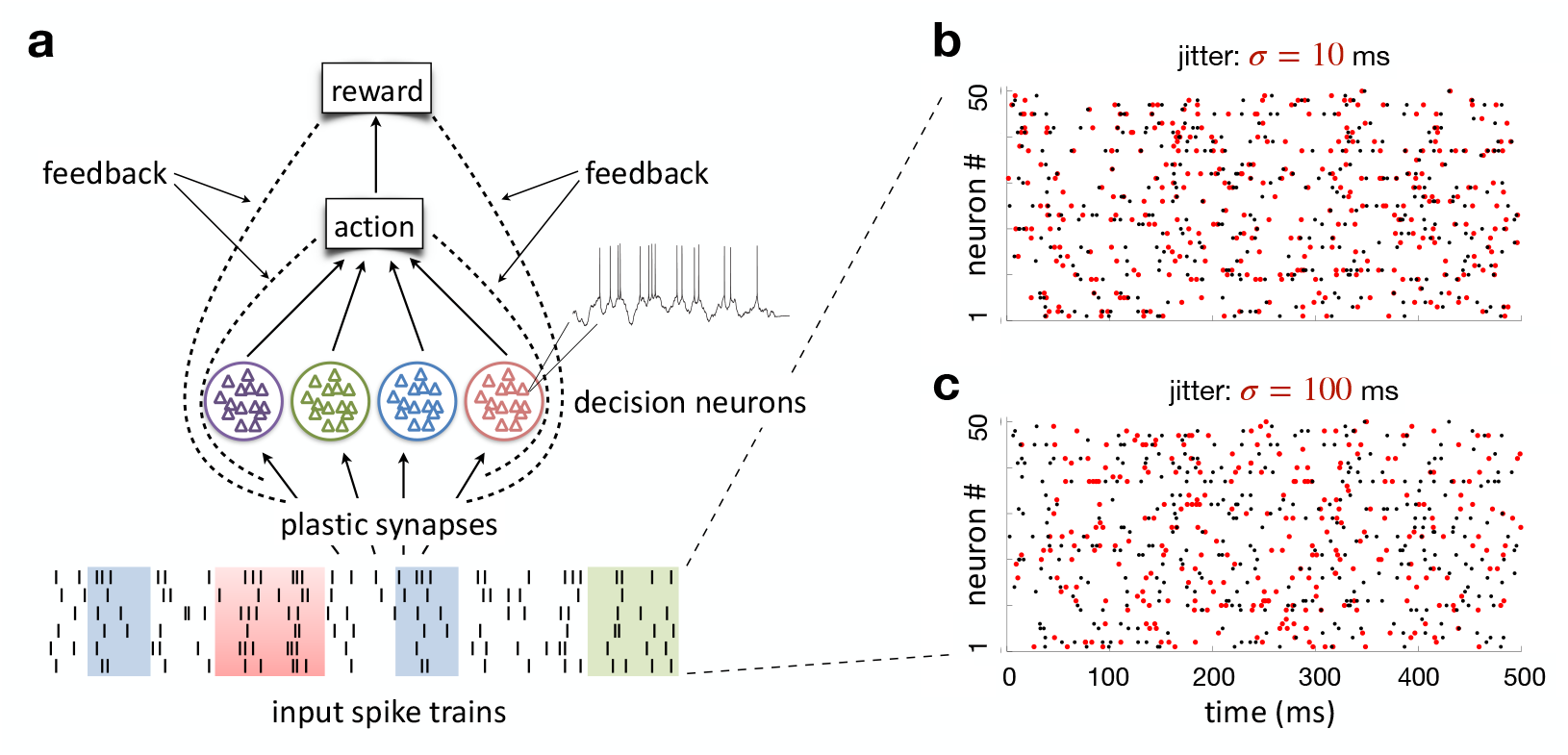
Stimuli and network architecture. **a)** Network architecture. Spike trains (‘input spike trains’) from *M* input neurons project to *N*_*dec*_ populations of spiking neurons (colored circles) through plastic synapses. Each population is responsible for a different decision, which occurs when a measure of the spike count rate of each population first crosses a predefined threshold. After each decision (and only then), feedback from reward (right vs. wrong) and the decision neurons’ activity determines the change of synaptic weights between the input neurons and the population neurons responsible for the current decision, according to Eq. 12. **b)** A segment of the input stream comprising 50 spike trains. Each line is a spike train and each dot is a spike time. The segment shown represents one stimulus prototype (black raster) together with a jittered version of it (red raster; 10 ms random dispersion around the original spike times, see Sec. IV D of Methods for details). The prototype could represent a relevant or a non-relevant segment of the stream. **c)** Same prototype shown in panel b), this time jittered by 100 ms (red).

The agent had to learn to detect the presence of a relevant stimulus by initiating an ‘action’, while ignoring the non-relevant segments (Fig. 1**a**). Only the appropriate response to relevant segments would lead to reward, while no reward was given for incorrect responses or for actions taken in the presence of non-relevant stimuli. An action occurred when the readout of the activity of one of the populations (a measure of its cumulative spike count rate) hit a fixed threshold. A correct decision was met with reward and the removal of the stimulus, as if a target had been hit or captured. Every action incurred a cost (a small negative reward), to discourage the agent from taking actions with high frequency. Neither reward nor cost was given in the absence of actions.

The agent learned to segment the input stream by trial and error, using a synaptic plasticity rule modulated by reward and population activity (Fig. 1**a**):

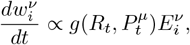

where 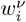 is the synaptic weight from input neuron *i* to decision neuron *ν* of population *µ, R*_*t*_ is the reward (or cost) received at time *t*, 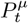 is the feedback on the spike count rate of population *µ, g*() is a function of *R*_*t*_ and 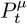, and 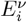 is the ‘eligibility trace’ of the synapse. The latter detects the covariance between the presynaptic spikes and the postsynaptic neural activity; only synapses with sufficient eligibility are modified. The detailed description of the plasticity rule and the network model is reported in Methods.

The plasticity rule reinforces correct decisions taken at the right time. It is a variation of a learning rule devised to work in more traditional tasks, where input stimuli are all relevant and feedback is given at the end of each stimulus presentation [6]. As mentioned in the Introduction, here we deal with a much more difficult problem, since the agent does not know what is relevant and feedback is obtained only in the presence of a decision. This problem is akin to temporal stimulus segmentation combined with a multiple-choice decision making task. As we show next, our learning rule can solve this problem in a variety of scenarios.

#### Detection task

We first illustrate the problem in the simplest possible scenario of one relevant stimulus embedded in an input stream of non-relevant ones. A single population of decision neurons (the learning agent) was trained to respond whenever the relevant stimulus was on. As shown in Fig. 2, the agent learned to successfully identify the relevant stimulus and respond within 50 ms (or less) from its onset, while learning to ignore the other stimuli (Fig. 2**b**). This was achieved despite using equal spike train statistics for relevant and non-relevant segments (notice how the input stream appears uniform on visual inspection, see e.g. the spike trains in black in Fig. 2**b**). The average learning curves across 150 training sets is shown in Fig. 3**a**. High performance on relevant stimuli was achieved at different times across independent training sets, depending on the initial weights, the input prototypes and stochastic decisions (dotplot in Fig. 3**a**). We call the time to achieve high performance the ‘learning threshold’, and quantify it as the first of 100 consecutive correct responses to relevant stimuli, taken as a criterion that learning has been accomplished. The learning thresholds were reached in every training set and were broadly distributed across independent training sets, as shown in the inset of Fig. 3**a**. Despite this variability, the average learning curve appears to improve gradually over time. Note that a variable learning threshold is an unavoidable consequence of the nature of the task: if the agent is initially set to take random and very sparse decisions, feedback is rare and so are synaptic changes. However, once the agent reaches the learning threshold for relevant stimuli, it quickly reaches the maximal performance for those stimuli. In the detection task, fewer than 250 presentations per stimulus were sufficient to reach criterion for half of the training sets (see inset of Fig. 3**a**).

**FIG. 2.**
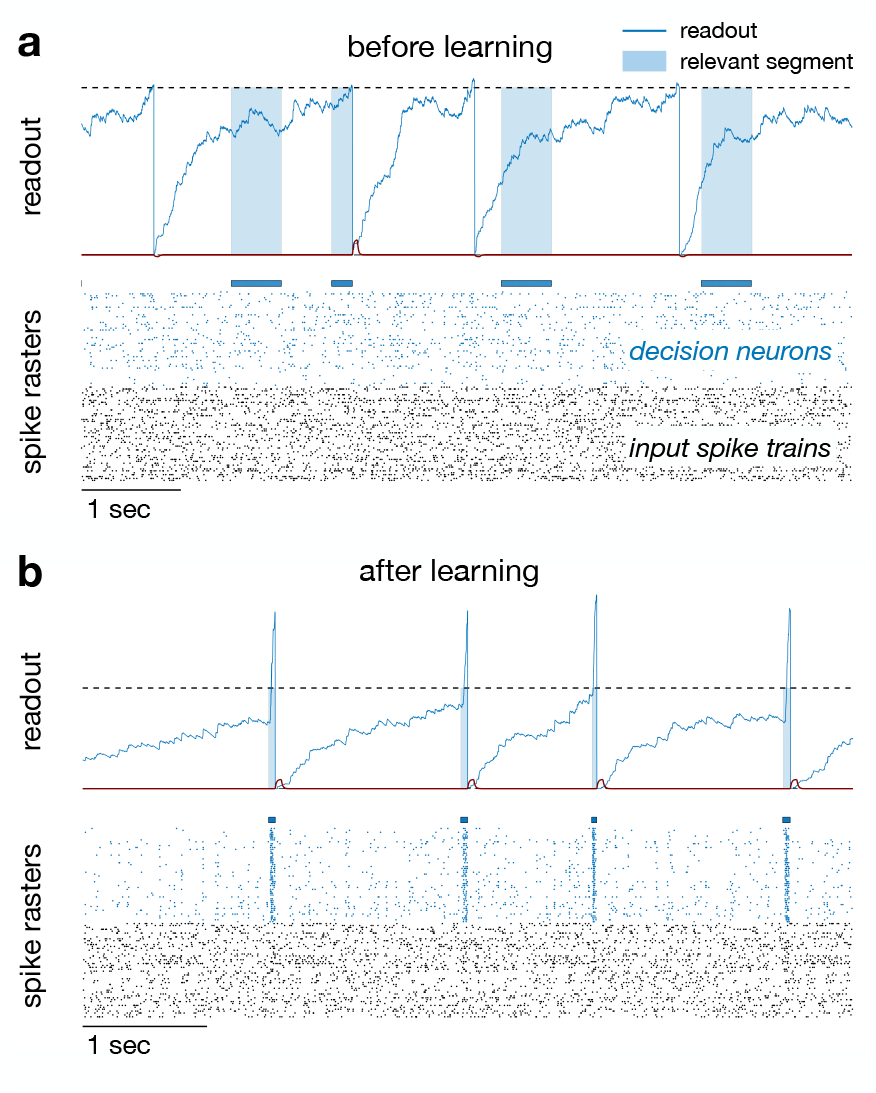
Learning the ‘detection’ task. **a)** Before learning. *Top panel:* readout of the single population of decision neurons, 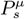 (blue, low pass filter of the population spike rate; see Eq. 2 of Methods). A decision was taken when 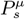 crossed a threshold Θ_*D*_ = 200 spikes/s (dashed horizontal line). Colored shadings above the rasters mark the presence of a relevant stimulus. Thirty milliseconds after a decision, the population readout was reset to zero. If the decision was a correct decision in the presence of a relevant stimulus, the stimulus was removed and reward was given (red line at the bottom). The initial decisions reflected the random initial weights of the synaptic connections between the input spike trains and the decision neurons. *Bottom panel:* raster plot of neural activity, with input spike trains in black and output spikes of decision neurons in blue. **b)** After training, the population readout correctly (and rapidly) reaches the threshold only in the presence of the relevant stimulus. In the input stream were embedded 4 prototypical segments acting as stimuli, of which 1 relevant (present during the blue shadings) and 3 non-relevant (present outside the blue shadings). Learning was assessed after 2,000 presentations per stimulus; *N* = 50 decision neurons and *N*_*i*_ = 50 input spike trains were used.

**FIG. 3.**
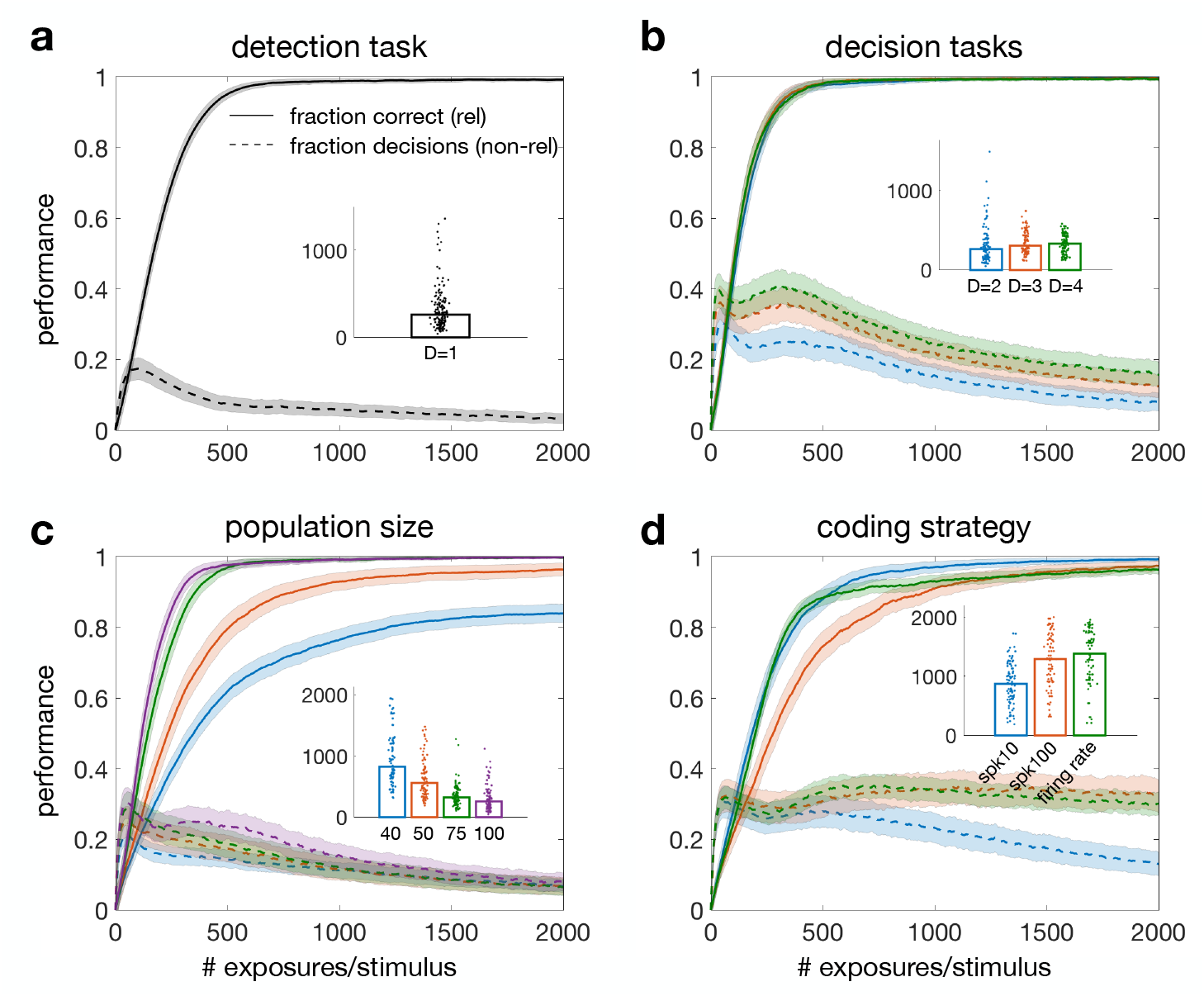
Average learning curves in temporal segmentation tasks. **a)** Detection task with 50 input spike trains and *N* = 50 decision neurons (same model and task as in Figure 2). *Solid curve:* average fraction of correct decisions with relevant stimuli across 150 training sets. *Dashed curve:* average fraction of (any) decisions in the presence of non-relevant stimuli (note: perfect performance with non-relevant stimuli is achieved when the dashed line converges to zero). **b)** Learning curves across 100 training sets for 3 different tasks having *D* = 2 (blue), 3 (red) or 4 (green) decision classes and one relevant stimulus per class out of 8, 12 and 16 total stimuli, respectively. In each case, there were *D* decision populations (regardless of the number of stimuli), each containing 100 (blue), 150 (red) and 200 (green) neurons. **c)** Increasing the number of decision neurons reduces the learning time. Shown are the mean learning curves for the binary decision task with 8 stimuli and *N* = 40, 50, 75 or 100 decision neurons per population (means across 100 training sets except for *N* = 40 where 200 sets were used). **d)** Learning with different encoding strategies: firing rate (green) and spike timing coding with a jitter *σ* = 10 ms (blue) and *σ* = 100 ms (red). The learning curves were averaged across 200 training sets for firing rate coding (green) and across 100 training sets in the remaining tasks. In all panels, shaded areas define SEM bounds across training sets. *Insets:* dotplots of learning thresholds (see the text) across training sets for each task (color coded as in the main plots); bars show median learning thresholds.

#### Multiple choice tasks

To implement multiple-choice decisions tasks, the decision neurons were partitioned in multiple populations (Fig. 1a). The different populations encoded different decisions, i.e., were responsible for learning stimuli belonging to different decision classes.

Fig. 3**b** shows successful performance in a segmentation task occurring simultaneously with a multiple-choice decision task with 2, 3 or 4 relevant stimuli (in the presence of 6, 9 and 12 non-relevant stimuli, respectively; learning thresholds were reached in every training set). An example of network activity before and after learning the 3-way decision task is shown in Fig. 4.

Note from Fig. 3**b** the nearly perfect overlap of the learning curves for the different tasks, achieved by increasing the number of decision neurons as the task gets more complex. This suggests that learning becomes faster when increasing the number of decision neurons [6], as confirmed in Fig. 3**c**. This panel shows the performance in the binary decision task when different numbers of decision neurons per population are used. As shown in the figure, a larger population size resulted in a faster learning speed (the learning rate was kept constant to allow for proper comparison).

Next, we tested whether the model could handle tasks with multiple relevant stimuli in each decision class, as well as various levels of stimulus sparsity (i.e., ratio of relevant to non-relevant stimuli; see Fig. S1). Although learning was slower for either a larger number of stimuli or for larger ratios, the average performance after training was high in all cases (ranging from 94% to 99% on relevant stimuli; see Fig. S1).

### B. Robustness to encoding strategies

Stimuli encoding into patterns of spike trains (Fig. 1**b-c**) was meant to reflect two widespread features of cortical activity: the relative stability of firing rates across trials coupled to unreliability in the precise spike times [7–9]. Learning in our model was very robust to jitter in the spike times (a proxy for trial-to-trial variability), tolerating amounts of jitter so large as to cover the entire duration of the stimulus (Fig. S2). The latter case was obtained by generating the spike times anew at each stimulus presentation and represents a full firing rate code.

The model was also robust to variations in coding strategies. We tested the model in a ‘spike timing’ coding mode wherein all input spike trains were generated from a Poisson process with the same firing rate. Although differences in the prototypes’ spike counts emerge from sampling, they are smaller than in the default stimuli, and therefore the information contained in the spike times is now much more relevant (in very long spike trains, the spike times would be the only way to discriminate the stimuli). Learning was successful also in this case (Fig. 3d), despite the presence of substantial temporal jitter in the spike times (100 ms jitter, Fig. 3d). Learning was slower with larger jitter, but the asymptotic performance was similar in all cases. The performance with small jitter was comparable to the performance with full firing rate coding (shown in green in Fig. 3d).

Next, we tested the model’s robustness to reduced stimulus dimensionality (i.e., smaller number of input spike trains). Learning was faster for higher-dimensional input patterns, but reached perfect asymptotic performance for as few as 25 input spike trains (Fig. S3). Similar results were found with sparse connectivity, i.e., after reducing the number of synapses projecting to the decision neurons (Fig. S4). In this case, each decision neuron received input from a randomly chosen subset of input spike trains. There was no difference in learning speed between full and 80% connectivity, and no appreciable degradation of final learning performance for levels of connectivity down to 30% (Fig. S4).

Finally, we tested the model with correlated input stimuli. In all cases shown so far, the spike trains were independently generated. Cortical spike trains however have some finite degree of pairwise or higher order correlations [10– which should result in more difficult discrimination. We first tested the model on spike trains with known pair-wise correlations. These spike trains were generated with the method of the dichotomized Gaussian model [18], which allows to generate Poisson spike trains with desired firing rates and pair-wise correlations *ρ* (examples are shown in Fig. S5). After 2,000 presentations per stimulus, the model had reached perfect performance on relevant stimuli for *ρ* = 0.1 and *ρ* = 0.3, and nearly perfect performance for *ρ* = 0.5 (Fig. S5d); mean performance with non-relevant stimuli was 94% or higher for any *ρ*.

### C. Segmentation of cortical data

The correlated stimuli of the previous section allow to quantify the learning performance as a function of a controlled amount of pair-wise correlations. The ultimate test, however, is the ability to learn real cortical data. To this purpose, we built prototypes of relevant and nonrelevant segments from input spike trains recorded simul- taneously from the gustatory cortex of awake behaving rats [19, 20] (see Methods for details). The rats had to press a lever to obtain one of 4 tastants (sucrose, citric acid, sodium chloride and quinine). In between trials, rats also received passive delivery of tastes at random times. Data collected before and after passive taste delivery were used to build the prototypes (see Methods for details). Ensembles of 5 to 9 single-units were recorded in each session, and units from all sessions were combined to obtain spike trains from 101 neurons (such patterns for two sucrose trials are shown in Fig. 5a). Of these, 50 neurons were randomly chosen to create prototype stimuli (Fig. 5b; different subsets of neurons were sampled for different training sets). This dataset was particularly appropriate for our purposes due to its long inter-trial intervals (a proxy for non-relevant stimuli) and because the neuronal activity had been recorded simultaneously from multiple single units. The learning problems were the same as with the surrogate Poisson spike trains described earlier. In the binary decision task, relevant stimuli were chosen to be the neural responses to two tastants (e.g., sucrose and citric acid); the responses to three different tastants were used in the 3-way decision task, and the responses to all four tastants were used in the 4-way decision task. For each tastant, the portion of the recordings immediately following stimulus delivery was taken as the relevant segment, whereas recordings end-ing 1s before stimulus delivery were taken as non-relevant segments (see Fig. 5a-b and Methods for details).

**FIG. 4.**
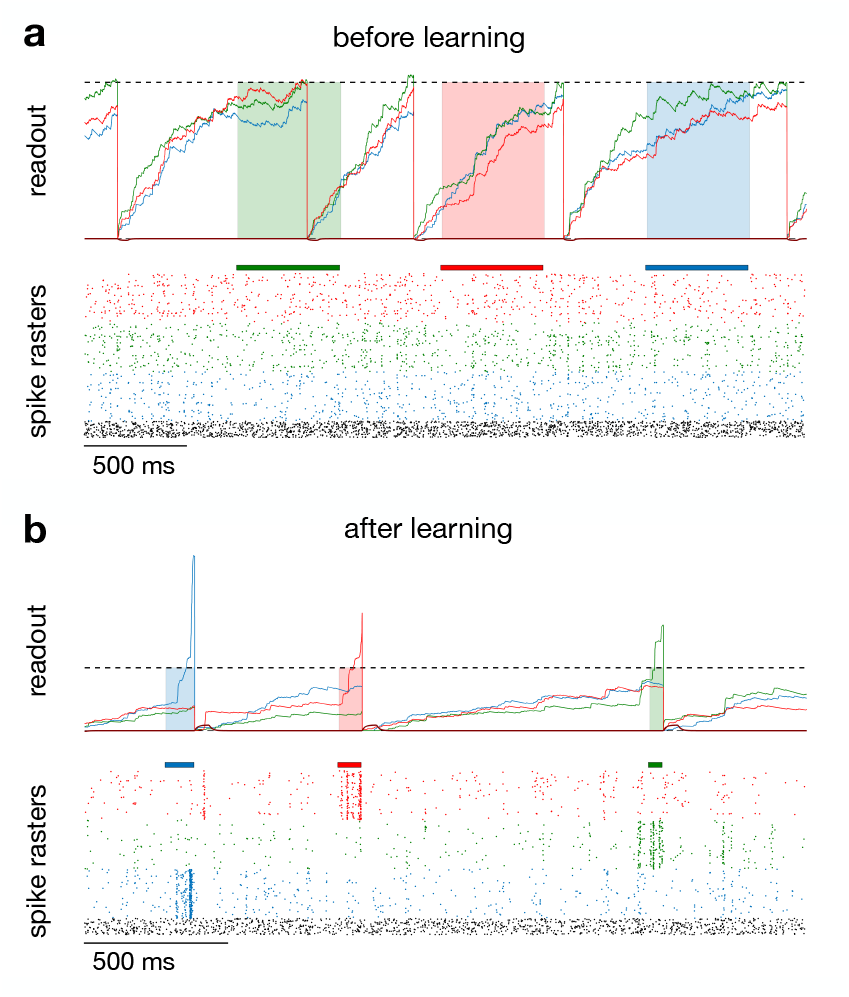
Learning a multiple choice segmentation task. **a)** Before learning. *Top panel:* readout of the population of decision neurons for three decision populations (color-coded; same keys as in Fig. 2). A decision was taken when the corresponding population readout crossed the threshold Θ_*D*_ (dashed horizontal line). Colored shadings above the rasters mark the presence of a relevant stimulus (different colors correspond to different stimulus classes). Reward was given after a correct decision in the presence of a relevant stimulus (see caption of Fig. 2 for details). *Bottom panel:* raster plot of neural activity, with input spike trains in black and output spikes of decision neurons in color (same color code as in top panel). **b)** After training. Same keys as in panel *a*. Learning was assessed after 2,000 presentations per stimulus; *N* = 150 decision neurons and *N*_*i*_ = 50 input spike trains were used.

**FIG. 5.**
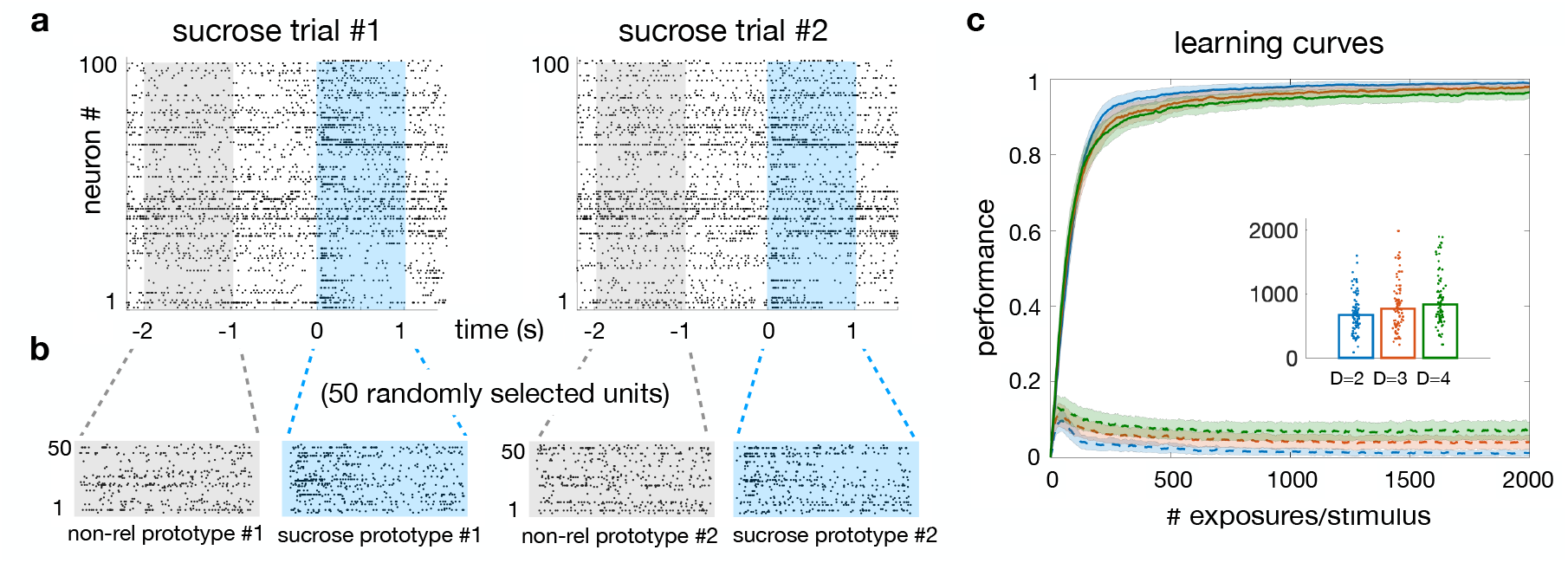
Segmentation of cortical data. **a)** Procedure used to build stimulus prototypes from cortical spike trains. Shown are spike trains from 101 neurons recorded over multiple sessions during the first two sucrose trials of each session (i.e., trials in which the tastant ‘sucrose’ was delivered). Taste delivery occurred at time zero. **b)** Generation of a single training set from cortical spike trains in (a). Data in a temporal window of [− 2, − 1] s were used to generate the prototypes for non-relevant stimuli (grey rasters); data in a temporal window of [0, 1] s were used to generate the prototypes for relevant stimuli (blue rasters). The same random subset of 50 neurons were used for all relevant and non-relevant stimuli in each training set. Prototypes for different tastants were built in the same way. Different random sets of neurons were used to generate different training sets (see Methods for details). **c)** Average learning curves with the stimuli obtained as shown in panels (a-b) for 3 different tasks having 2 (blue), 3 (red) or 4 (green) decision classes and 1 relevant stimulus per class out of 8, 12 and 16 total stimuli, respectively. Each curve is the average across 100 training sets generated from cortical data as described above (same keys as in Fig. 3b). *Inset:* dotplots of learning thresholds for each task. Learning thresholds were reached in 87% (*D* = 4), 92% (*D* = 3) and 100% (*D* = 2) of training sets.

The results are shown in Fig. 5c and are similar to the results obtained with surrogate spike trains (Fig. 3b). We also tested the ability of the trained agent to generalize to unseen stimuli of the same class (i.e., elicited by the same tastant, but in trials not used for training). The results are reported in Table S1 (column ‘generalization’). To assess the quality of generalization, its performance was compared to the performance on unseen stimuli of a different class (i.e., elicited by a different tastant than used for training; see Methods for details). As shown in Table S1 (compare columns ‘generalization’ with ‘control’), the generalization performance was significantly higher than control in all tasks (*p <* 0.015 or lower, Mann-Witney U test; see the caption of Table S1 for details).

Note that the low absolute value of generalization performance (*<* 50%) was the consequence of training on one stimulus per class while testing on 6 unseen stimuli of each class. Generalization performance can be boosted by the common practice of training on multiple stimuli. For example, training in the binary decision task with 6 stimuli per class, and then testing on one unseen stimulus of each class, gave a mean generalization performance of 74% (last row of Table S1).

## III. DISCUSSION

Learning to abstract relevant information from the environment is a crucial component of decision making; yet, standard approaches typically assume that the relevant stimuli, and especially the times at which they occur, are known to the decision maker [21–23]. Even assuming partial ignorance about the current state or about the rules of a decision task [1], there is usually no ambiguity about the presence of a cue that, possibly incompletely, signals the current state. This is especially true for the type of decision tasks typically modeled by signal detection theory [24]. For example, in psychophysical discrimination tasks with random dot displays [25–27], the aggregate direction of motion may not be perceptually clear, but the presence of the random dot display occurs at known times and there is no doubt on its relevance to the task.

In the problems studied here, relevant stimuli are continuously embedded in spatiotemporal patterns of neural activity, and occur at times that are unknown to the agent. Such a task requires the temporal segmentation of the input stream, which is possible only if the agent takes the correct action at the right time. Thus, both the presence of relevant segments and the correct action associated to each segment must be discovered by the learning agent. Since this learning process ends up assigning meaning to the relevant segments, it is tempting to speculate that this reward-based mechanism of parsing input streams may be one way to form concepts out of seemingly unstructured stimuli [28, 29].

Learning in this scenario was accomplished in a biologically plausible model with populations of spiking neurons. The model is plausible in several ways: i) it uses a local learning rule based on preand postsynaptic spikes as well as post-synaptic potentials, all quantities directly accessible to each synapse; ii) the learning rule receives neuromodulatory feedback from reward and decisions [30, 31]; (iii) learning can be achieved with erratic cortical spike trains using different coding strategies (spike timings or firing rates), in the presence of variable degrees of correlations, and with no need to customize the learning rule to each case separately. In the following, we discuss the salient features of our approach and later compare it with alternative approaches for reinforcement learning and decision making.

### A. Scaling performance with population size

Reinforcement learning with spiking neurons is notoriously slow (e.g. [32–34]) and does not scale well with population size [35–37]. This is because larger populations of neurons make the spatial credit assignment problem more severe, resulting in slower learning [6, 38]. A slowdown of learning speed with population size is also contrary to the principle of population coding, according to which performance in any task should increase with population size [39]. Thanks to a learning rule that averages the gradient of the reward over the outcomes of each neuron, in our model learning speed increases with the size of decision populations (Fig. 3**c**). This means that increased task difficulty can be compensated for by increasing the number of decision neurons, so that similar learning speeds can be reached in more difficult tasks, for example, when increasing the number of decision classes (Fig. 3**b**), the number of input stimuli, or the number of non-relevant stimuli embedded in the input stream (Fig. S1).

### B. Flexibility to coding schemes

Our learning rule is also robust to a large class of coding schemes and can handle real cortical datasets. We have shown successful learning in 3 different coding scenarios: i) when stimuli are coded by prototypical spike patterns, despite large amounts of jitter in the spike times (Fig. 3**b** and Fig. S2), ii) when stimuli vary only in their spike timings across neurons, with equal firing rates across neurons (spike timing coding, Fig. 3**d**)), and iii) when stimuli vary both in their firing rate patterns and in their single neurons’ spike times (a full firing rate code; green curve in Fig. 3**d**). We emphasize that in the case of precise spike patterns (scenario ii), all stimuli had exactly the same statistics, and thus they could not be encoded as firing rate vectors, nor could be learned by an unsupervised algorithm that looks for idiosyncratic regularities in the stimuli. Successful learning occurred also in the case of stimuli encoded as firing rate patterns, where the spike trains had different firing rates but were generated anew at each stimulus presentation. In these more challenging tasks learning was correspondingly slower, albeit still comparable to the main coding scenario, as shown in Fig. 3**d**. Moreover, the model performs well on stimuli with correlated spike trains (Fig. S5) as well as cortical spike patterns recorded from behaving animals (Fig. 5; more on this later).

The fact that learning is successful in the case of lax spike timing reproducibility is particularly appealing, giving that cortical spike trains are notoriously erratic. Our results suggest that our model could work in the face of the large trial-to-trial variability observed in cortex [40, 41]. This is notable given that the learning rule was initially designed to handle spike patterns that are precisely timed. We note that, within a framework similar to ours, learning rules that depend directly on firing rates can be built, an approach that would allow for faster learning [42], at least in more traditional tasks. We decided not to use this approach for the segmentation task of this paper, in part for reasons of biological plausibility, as it is not clear how plasticity could depend on firing rates directly; and in part to obtain a more general learning rule that could handle a wider range of codes. The problem of neural coding is an open question and this approach allowed us not to commit to a specific coding scenario. Our results demonstrate the flexibility of our model in dealing with coding schemes interpolating between precise spike timing and firing rate coding with various degree of pairwise correlations, while also showing robustness to low levels of connectivity (Fig. S4) and stimulus dimensionality (Fig. S3).

Regarding the decision code, we have used the simple and widely used approach of having separate populations of neurons competing for decisions [2, 43]. Although it is possible, in principle, to adapt the model to handle decisions based on other coding scenarios, such as phase coding [44–46] or latency coding [38, 47], we have not tested these possibilities. The emphasis of this work was on learning to segment unknown stimuli, with less relevance given to the decision mechanism. It would be of interest to develop a biologically plausible model capable of learning its own ‘valid’ actions in addition to its own relevant stimuli. This is left for future work.

### C. Learning cortical spike patterns

The model can segment cortical spike train patterns recorded in the gustatory cortex of behaving rats, with good generalization performance to unseen segments recorded under the same conditions. Testing the model on cortical data has two main advantages over surrogate stimuli. First, it allows a fairer evaluation of the generalization performance. We expect generalization to surrogate stimuli of the same class to be easy, given that unseen stimuli would be generated by the same prototypes used for training. Secondly, it allows to quantify the ability of the model to extract essential features of the data related to the external stimuli. This is particularly relevant during generalization. Evoked cortical patterns in the gustatory cortex can be successfully decoded [48, 49], suggesting that some common features of the data are preserved across stimulus presentations. However, it is not clear how to extract these common features. Previous efforts have analyzed the higher order correlations of cortical spike trains in search for identifiable features [11–15, 50], however the analysis of high-order correlations and their relevance to pattern features is a difficult task. It is important, and somewhat reassuring, that we can learn to extract the essential features of spike patterns by reinforcement of correct decisions.

### D. Precision vs. recall

The performance of the model on relevant stimuli was generally higher than on non-relevant stimuli, especially with surrogate data. The tasks themselves were not symmetrical, since non-relevant stimuli must be simply ignored and produce no reward. It is possible to improve performance with non-relevant stimuli at the expense of performance with relevant stimuli by choosing a different cost/reward ratio. Our main goal, however, was to demonstrate the ability to segment unknown relevant stimuli. In order to learn, the agent must explore by taking random actions, and acting during non-relevant segments is a reasonable price to pay to discover the relevant ones. In the language of pattern recognition, this issue can be stated in terms of ‘precision’, the fraction of relevant stimuli among those detected as relevant (i.e., true positives/(true positives+false positives), vs. ‘recall’, the fraction of relevant stimuli identified as such (i.e., true positives/# of positives) [51, 52]. Lower performance with non-relevant stimuli lowers the precision while keeping the recall intact. We have basically prioritized the recall, i.e., the performance on relevant stimuli, which is close to perfect in all our tasks. As learning problems become more difficult, it is reassuring to know that we may still attain a high recall at the cost of losing precision, if necessary.

### E. Comparison with previous work

Segmentation tasks are often solved by unsupervised methods such as Hidden Markov Models (HMMs) [53], which have been successfully applied to the segmentation of cortical patterns of spike trains similar to those modeled here [19, 47, 48, 54–56]. However, HMMs require a-priori knowledge of the structure of the relevant stimuli (e.g., Poisson-HMM vs. Gaussian-HMM) and they also require knowledge of the total number of relevant stimuli (or hidden states, in HMM parlance). Although non-parametric variations of HMM exist [57, 58], they are based on recursive offline algorithms. Online procedures to approximate the expectation-maximization steps required in HMM can learn to perform tasks based on the temporal statistics of stimuli [59–62], but would not be able to learn to segment a stream where the input spike trains all have the same underlying firing rate, as is the case of the spike timing code of Fig. 3**d**.

Our model also differs from the class of recurrent neural circuit models of decision making [2]. As most traditional models of decision making, these models require a neural population encoding each input stimulus, whose identity and timing are consequently known.

Similarly, in previous models of reward-modulated learning by spiking neurons in continuous time [34, 35, 42, 63, 64], the relevant stimuli are known and need not be inferred via segmentation of the input stream.

Previous work on detecting spatio-temporal spike pat-terns in continuous streams [65, 66], a task closer to ours, has mostly used unsupervised learning, especially through spike-timing-dependent plasticity (STDP). In these approaches, the main focus has been on segmentation and representation rather than decisions; however, it is possible that appropriate actions could be learned by a separate system. For example, in Ref. [65] the authors have used STDP to train a spiking network to produce stimulus-specific firing patterns under a variety of conditions. In this case, decisions could be learned by a separate reinforcement learning system trained on the stimulus-specific patterns. Since STDP is very sensitive to spike-timing jitter [67], these methods have used ‘frozen’ spike patterns or only very limited spike timing jitter in the stimuli, and presumably cannot handle firing rate patterns.

Perhaps closest to our problem and approach is “aggregate-label” learning [68]. In this approach, a neuron or a small network are trained to respond to different stimuli with a different number of spikes to each, with no *a priori* knowledge of which stimuli are relevant and when they are being presented. Unlike our model, aggregate-label learning is supervised or self-supervised, using feedback on the total number of spikes required in the current trial. This requirement, and the learning rule itself, seem less biologically plausible than our model. On the other hand, aggregate-label learning can solve a non-Markovian task, because the feedback is only given at the end of a trial containing many stimulus presentations. This is currently beyond the reach of our model, which perhaps could be extended along the lines of Ref. [69], as discussed below.

### F. Extensions

Our work can be extended in a number of directions. One possible direction is to include non-stationary stimuli. In one example, spike patterns slowly appearing and disappearing on top of a noisy background would have to be detected and segmented according to action-relevance. Different subsets of sensory neurons would encode different stimuli, while the remaining neurons would represent streaming noise (different subsets for different stimuli). Preliminary computer experiments show encouraging results, at least in the presence of 2-8 stimuli with negligible coding overlap (not shown). Another possible direction is the extension to sequential decision tasks [69], wherein actions and/or rewards are required only after a sequence of relevant segments. In such a scenario, decisions depend not just on current state but also on previous history, and proper segmentation could lead to context-dependent representations of relevant stimuli (i.e., according to the sequence in which they are embedded [70]). Neural representations of planning sequential behavior are essential to many complex behaviors and are related to concept learning and the representation of abstract rules [5, 28, 71–75]. An extension of our framework to deal with more complex embeddings of stimuli could potentially lead to context-dependent representations of relevant events or even abstract concepts. A comprehensive investigation of these directions is left for future studies.

## IV. METHODS

### A. The network model

#### Task and network architecture

The model network comprised *N*_*dec*_ populations of spiking neurons receiving input spike trains via plastic synapses (Fig. 1A), with *N*_*dec*_ varying from 1 to 4 (see Table S2 for all parameter values). Unless stated otherwise, each decision population comprised *N* = 50, 100, 150 or 200 neurons in tasks with *N*_*dec*_ = 1, 2, 3 or 4 decision classes, respectively, and received *N*_*i*_ = 50 input spike trains. Except in the study of reduced connectivity (Fig. S4), all input spike trains projected to all decisions neurons. Each population *µ* coded for a specific decision and initiated the corresponding action whenever a running measure of population spike rate 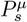 (defined below) crossed a decision threshold Θ_*D*_ = 200 spikes/s (the same for all populations). As long as all population scores were below threshold, no decisions were taken and no feedback was given to the agent.

Stimuli were continuously presented to the agent as streaming spatio-temporal patterns of spike trains, and were divided into relevant and non-relevant stimuli (details in Section “Stimuli and computer experiments” below). Each relevant stimulus was associated with a correct decision, one among the *N*_*dec*_ possible decisions. Thirty millisecond after a decision occurred, the stimulus was removed and rewarding feedback was given. Every decision (whether correct or incorrect) incurred a small cost (*R* = ™0.1) to deter the agent from continuously taking actions; positive reward (*R* = 1) was given only for a correct decision in the presence of a relevant stimulus (netting a total reward of *R* = 0.9). In the (rare) case of multiple populations reaching threshold simultaneously, only one was selected pseudorandomly as the population responsible for the decision.

The model and simulation parameters are summarized in Table S2.

#### Population scores and decision dynamics

A natural measure of ongoing population activity is the spike count rate,

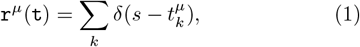

where 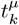 are the times of the spikes emitted up to time *t* by neurons of population *µ*, and *δ*(*t*) is Dirac’s delta function. To compute this quantity during the presentation of a stimulus, the learning agent would need to know when a stimulus starts and ends. To avoid this, `r`^*µ*^(`t`) was approximated by a low-pass filter

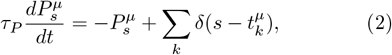

with time constant *τ*_*P*_ = 500 ms matched to the stimulus durations (different *τ*_*P*_ values between 50 and 2, 000 ms affected the learning curves but did not prevent learning). Eq. 2 defines the online ‘population score’ 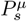, which is responsible for initiating a decision. A decision was said to have occurred at the first time *t*_*d*_ such that 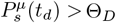 for some population *µ*, and at time 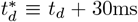 all population scores 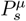 were reset to zero. The 30 ms delay between the decision and the reset adds biological realism while allowing a confidence signal to build up, as follows. At time 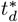 following a decision by population *µ*, a feedback on the decision is given to all neurons in the population through the quantity 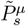 obeying

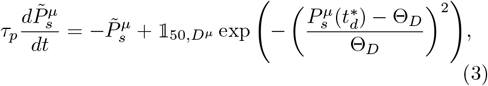

with *τ*_*p*_ = 50 ms and 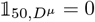 except for 50 ms after 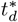, when it is set to either +1 or to 1, depending on whether or not the population *µ* was responsible for the current decision. The driving term 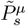 has a double role: its magnitude can represent the population’s own confidence in the decision at time 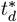, and as such it was used to attenuate learning (details in Sec. IV C); its sign was used to build an individualized reward signal as proposed in [6] (see Eq. 13).

### B. The neuron model

The decision neurons producing scores 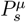 were modeled as spiking neuron models driven by post-synaptic potentials and having a probabilistic threshold for spike emission [76]. Specifically, neuron *ν* was defined by membrane potential *u*^*ν*^ obeying

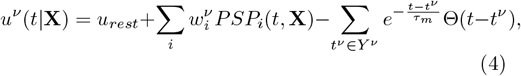

where: *u*_*rest*_ = − 1 is the resting potential (in arbitrary units); 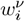 is the synaptic weight connecting neuron *ν* to its presynaptic neuron *i*; **X** *X*_1_, *X*_2_, …, *X*_*M*_ is the collection of the *M* presynaptic input spike trains, where 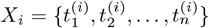 are the spike times of the *i*^*th*^ spike train; *PSP*_*i*_(*t*, **X**) is the postsynaptic potential due to the input spike train *X*_*i*_,

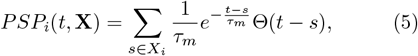

with *τ*_*m*_ = 10 ms being the membrane time constant of the neuron; *Y* ^*ν*^ is the output spike train (comprising spike times *t*^*ν*^); and Θ() is the Heaviside function (Θ(*t*) = 1 for *t >* 0, and Θ(*t*) = 0 otherwise). A neuron with membrane potential *u* in a short time bin *dt* emitted a spike with probability

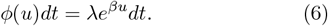

Here, *β* = 5 controls the stochasticity of spike emission whereas *λ*, the firing rate at *u* = 0, has an additive or subtractive effect on spike probability, i.e., changes in *λ* shift *ϕ*(*u*) rightwards or leftwards. *λ* was slowly modified during learning to help keep the decision populations’ activities near the decision threshold (see Eq. 15).

Given input spike pattern **X** lasting from 0 to *t*, the log-likelihood that a decision neuron produces the output spike train *Y* ^*ν*^ is [76]

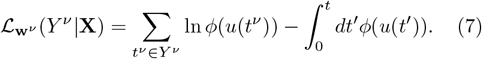

### C. The learning rule

The learning rule was designed to maximize a measure of reward during the fully online learning scenario described in the main text. The starting point is the online learning rule performing stochastic gradient ascent in the average reward, which requires the temporal derivative of the gradient of the log-likelihood 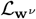 [6]:

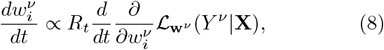

where *R*_*t*_ is the reward at time *t*. From Eqs. 4, 6, 7, we get

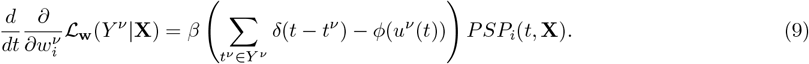

Since the synapses do not know the onset and offset times of the stimuli, we replace the temporal derivative of the gradient with a low-pass filter,

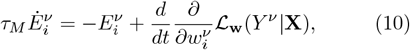

where *τ*_*M*_ = 500 ms. 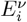 is called ‘eligibility trace’ of synapse 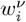.

With these quantities in hand, an online learning rule performing gradient ascent in the average reward would read

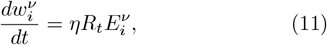

where *η* is the learning rate and *R*_*t*_ is the reward at time *t*. However, to alleviate the spatial credit assignment problem, we follow [6] and replace the global reward with an individualized reward signal *r*^*ν*^ that equals 1 if neuron *ν* took the right decision, and equals 1 otherwise (see subsection ‘Individualized reward’ below for its biological implementation). Specifically, we used the learning rule

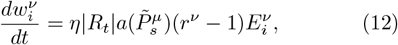

for each synapse onto postsynaptic neuron *ν* belonging to population *µ* responsible for the current decision. Although this learning rule can be applied to all synapses, regardless of whether or not their post-synaptic neurons were responsible for the decision, we found it sufficient to apply it only to the latter. The factor 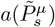 is an attenuation factor equal to 1 (no attenuation) for an incorrect decision (*R*_*t*_ *<* 0) and equal to 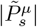 for a correct decision (*R*_*t*_ *>* 0), with 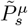 given by Eq. 3. Since *R*_*t*_ is different from zero only around a decision time (when reward is given; see subsection ‘Individualized reward’ below), the factor *R*_*t*_ confines the synaptic update to a temporal window around reward delivery. In all numerical experiments reported here, *η* = 10 ms (note that 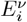 has units of 1/ms^2^).

Before describing each term of the learning rule in detail, we make two observations:

i. In the case of episodic learning with spike/no spike coding, where decisions are binary and based on a majority rule in a single population, the learning rule Eq. 12 performs stochastic gradient ascent in a monotonic function of reward and population activity, as proved in [6]. The need to introduce the individualized reward signal *r*^*ν*^ in this rule arises because otherwise learning worsens as the population size increases [6]. Instead, learning through Eq. 12 speeds up with population size, as demonstrated in [6] for more standard tasks and in this work (Fig. 3c) for the combined segmentation/decision-making task.
ii. The right hand side of Eq. 9 contains the product of a postsynaptic term (the difference between the actual firing rate 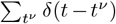 and the average firing rate *ϕ*(*u*^*ν*^) of neuron *ν*), and a term reflecting presynaptic input (the postsynaptic potential *PSP*_*i*_). Together with the reward-dependent term in Eq. 12, the latter is a three-factor synaptic plasticity rule as found e.g. in cortico-striatal synapses, a major locus for reinforcement learning [77].

#### Learning attenuation

According to Eq. 12, only the synapses targeting neurons voting for the wrong decision (*r*^*ν*^ = ™1) are updated after a decision. The decision itself may be correct or incorrect. The update is full in case of an incorrect decision and attenuated by a factor 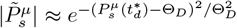 (see Eq. 3) in case of a correct decision. The rationale for this choice is the following: if 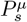 is large or small compared to Θ_*D*_ soon after a *correct* decision, the population is highly confident of its vote and no further learning is required (and indeed in this case 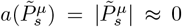 in Eq. 12); otherwise, if 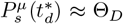, the population is maximally undecided about the current decision, and full learning is required 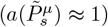.

#### Individualized reward

The individualized reward signal could be made available locally at each synapse by broadcasting feedback from the population activity via three neuromodulatory signals: 1) 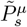 (representing the deviation from equilibrium of the concentration of a neurotransmitter such as acetylcholine or serotonin), 2) each neuron’s own activity 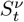 (e.g., through intracellular calcium transients), and 3) the global reward feedback *R*_*t*_ (e.g., through extracellular dopamine). Using these three signals, an online estimate of the individualized reward signal for the *ν*^*th*^ neuron in population *µ*, to be used in Eq. 12, was computed as

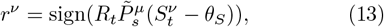

where *θ*_*S*_ = *e*^*−*1.1^ is a threshold for 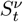 (see below). The reward signal *R*_*t*_ was a low-pass filter of the global reward *R*, transiently driven by *R* for 50 ms after each decision time 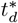, and then decaying to zero exponentially with time constant of 10 ms. 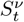 tracks the neuron’s spiking activity and was given by

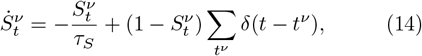

where the {*t*^*ν*^} are the neuron’s output spike times, and *τ*_*S*_ = 500 ms. If, for example, the decision was correct (*R*_*t*_ *>* 0) and population *µ* was above threshold 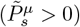, the neurons of the population which had been sufficiently active (*S*^*ν*^ *> θ*_*S*_) had voted for the correct decision, hence for them *r*^*ν*^ = 1, whereas those neurons that had not been active (*S*^*ν*^ *< θ*_*S*_) had voted against the correct decision, and for them *r*^*ν*^ = −1. One can similarly work out all other possible scenarios and confirm that the Eq. 13 indeed provides the correct feedback in each case. In the simpler case where the learning rule is applied only to synapses onto neurons responsible for the decision (as done here), 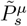 is always positive and individualized reward is driven by the covariation of reward and the posy-synaptic neuron’s own activity.

The parameters *τ*_*S*_ and *θ*_*S*_ were chosen with the following rationale [6]. If the time constant *τ*_*S*_ is equal to stimulus duration, then one or more spikes occurring in response to a stimulus yield a memory trace satisfying 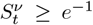 at stimulus ending. Since reward *R*_*t*_ peaks some short time later, we used a smaller *θ*_*S*_ = *e*^*−*1.1^.

#### Homeostatic mechanism

Neurons’ activities were kept close to the decision threshold via a homeostatic mechanism. This was implemented via the update of the parameter *λ* in Eq. 6 controlling the neurons’ spike probabilities. Specifically, for every neuron in population *µ, λ ≡ λ*^*µ*^ was changed according to

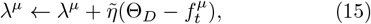

where 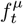 is a running average of the population spike count rate similar to 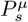 (Eq. 2) but without post-decision reset and a with much longer time constant (4 seconds). The update rate 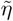 was 0.01 in all computer experiments. The update Eq. 15 occurred at random intervals distributed according to a uniform distribution between zero and 2 seconds. In between updates, *λ*^*µ*^ was kept fixed not to interfere with learning the weights. *λ*^*µ*^ was constrained to be positive and was the same for all neurons in population *µ*. Other update schedules gave similar results and in general the mechanism was very robust to the mean length of the interval between updates or the interval length distribution, see Fig. S6. Mean inter-update times as short as 50 ms also led to successful learning (Fig. S6). Deterministic schedules wherein *λ*^*µ*^ was updated at regular intervals (or every fixed number of presentations) worked just as well, however they require knowledge of the time (or the number of stimulus presentations) elapsed since the last update, and were dismissed as less biologically plausible. This homeostatic mechanism is beneficial because, as learning modifies the weights, it also induces changes in population activity. These may occasionally produce very low activity in some populations, resulting in lack of decisions for those populations. When this occurs, the agent is trapped in a local maximum of the average reward, and learning practically stops for that population. In such a case, the homeostatic mechanism slowly increases the firing probability in single neurons until the desired population activity level is restored and decisions are again possible.

### D. Stimuli and computer experiments

#### Main stimuli

The stimuli were jittered versions of prototypical spatiotemporal patterns of 50 Poisson spike trains with constant firing rates sampled from a uniform distribution between 2 and 24 spikes/s. In Fig. S3, the number of spike trains was varied from 25 to 200. Note that the choice of a Poisson process is convenient but not strictly necessary, i.e., other distributions could have been used instead, including the empirical distribution of cortical data [78, 79]. With the exception of Fig. S4, all-to-all connectivity between input spike trains and decision neurons was used. Each stimulus presentation lasted for an average 500 ms (with small variations due to spike times jittering, see below), except when removed early as a consequence of a decision (learning was robust to variations in stimulus durations). Stimuli prototypes were initially generated and then, prior to each presentation, temporal jitter was added to the spike times to produce a noisy version of the prototypes. For each spike time *t*_*sp*_, jitter was added by sampling from a Gaussian distribution centered on *t*_*sp*_ and having a standard deviation of *σ* = 10 ms (Fig. 1**b**), unless specified otherwise to test for robustness to larger values of jitter (Fig. S2). Stimuli were presented as a continuous input stream while learning occurred online as described in previous sections. The majority of the stimuli (a 3:1 ratio) were always defined as ‘non-relevant’ and the agent had to learn to ignore them not to incur the small cost attached to any decision (whether correct or incorrect; see Sec. IV A). Different ratios of relevant to non-relevant stimuli were used to test robustness as shown in Fig. S1. Contrary to standard reinforcement learning methods, decisions were not enforced at stimulus offset. Thus, in the absence of decisions, no learning would be possible.

#### Firing rate and spike-timing coding

For the results presented in Fig. 3d, stimuli were encoded by either firing rates only, or by spike timing only. In the former case, the prototypes were generated as for the main stimuli, but at each new presentation the spike trains were generated anew. This is akin to having a jitter in the spike times covering the entire duration of the stimuli. In spike-timing coding, the spikes trains of the prototypes had all the same firing rate of 12 spikes/s; the spike times were then jittered in time by *σ* = 10 or 100 ms before each new presentation.

#### Stimuli with specified pair-wise correlations

Spike trains with desired firing rates and pair-wise correlations were generated by using the dichotomized Gaussian model [18]. The same distribution of firing rates as for the main stimuli was used, with a desired pairwise correlation of 0.1, 0.3 or 0.5 (see Fig. S5 for examples). One hundred training sets were generated for each correlation level, and training proceeded as for the main stimuli. In particular, each presented stimulus was a jittered version of its prototype, with temporal jitter of *σ* = 10 ms.

#### Learning stimuli from cortical datasets

Cortical data were obtained from multi-electrode recordings of single units as described in [20]. Briefly, Long-Evans rats were trained to press a lever within two seconds of hearing a pure tone to obtain one of 4 tastes delivered directly into their mouth through an intra-oral cannula (this procedure allows high temporal resolution on stimulus delivery). Tastes and concentrations were: 100 mM sodium chloride, 100 mM sucrose, 100 mM citric acid, and 1 mM quinine. Water (50 *µl*) was delivered to rinse the mouth 5 seconds after the delivery of each tastant. Unexpectedly to the rats, the same tastants would also be passively delivered at random times during the experiment. In each session, rats performed 7 trials with passively delivered stimuli, and 7 trials with expected stimuli, for a total of 28 passively delivered tastants and 28 expected tastants. Multiple single-unit recordings from the primary gustatory cortex were collected while the rats performed the task. Neurons with firing rates lower than 2 spikes/s and neurons exhibiting a large peak around the 6-10 Hz in the power spectrum of their spike trains, potentially reflecting the motor response [49, 80, 81], were excluded. The remaining 101 neurons would form the patterns shown in Fig. 5a and were used to generate the stimulus prototypes as follows.

To obtain each training set, we randomly sampled 50 (different) input neurons out of the 101 cortical neurons (see Fig. 5b). To build relevant stimuli, we took the data falling in a temporal window between zero and 1 s after the passive delivery of a tastant; for non relevant stimuli, we took data falling in a 1 s window ending 1 second before the passive delivery of a tastant (during the so-called ‘ongoing activity’, [19]). Due to long and random intertrial intervals following water rinse (averaging about 40s), it is reasonable to assume that no residual trace of the previous stimulus was present at the time of a new delivery. Hence, the data prior to stimulus delivery can be considered non-stimulus related and therefore suitable to represent non-relevant stimuli. As with the main stimuli, cortical stimuli had average duration of 500 ms, and their spike times were jittered by 10 ms before each presentation. For each stimulus in each training set, we had 7 prototypes, corresponding to the 7 experimental trials with a given tastant; the 7 prototypes were considered as relevant stimuli of the same class (i.e., demanding the same decision; two prototypes for sucrose are shown in Fig. 5b). In each decision task of Fig. 5c, we used 1 relevant segment for each class (say, sucrose and citric acid from trial 1 for the binary decision task), and 3 times as many non-relevant segments, built as described above and randomly selected from the pool of all non-relevant segments. We trained the model on 100 training sets; for each new training set, we resampled the 50 input neurons from the 101 cortical neurons and repeated the procedure just described.

After training, we tested the ability of the model to generalize to unseen stimuli of the same class. Specifically, for each training set, we had the agent perform the same task, with plasticity switched off, using, as relevant stimuli, unseen stimuli from trials 2 to 7 of the same class. For example, if the training stimulus was in the class of sucrose, we tested the model on sucrose prototypes not used during training. These relevant segments were pitted against unseen instances of non-relevant segments. Each generalization test included 10 presentations per stimulus. In the binary task with 6 relevant stimuli per class (see the last row in Table S1), generalization was tested on the single, remaining unseen stimulus, of each class.

To establish the significance of the generalization performance, we compared the latter against the performance in a ‘control’ task where we used stimuli from a different class than the class used for training. For example, if sucrose and citric acid had been used for training in the binary decision task, sodium chloride and quinine were used for testing. For the 4-way decision task, in which all classes had been used for training, unseen instances of class 2, 3 and 4 were used as instances of class 1; unseen stimuli of class 1, 3 and 4 were used for class 2, and so on. Note that not only these stimuli had not been used during training, but they had been obtained in response to different tastants during the experiments. Therefore, performance in the control task was taken as a proxy for chance performance on unseen stimuli. The generalization performance across 100 validation sets (one for each training set) was compared with the performance in the corresponding control task using a Mann-Whitney U test (Table S1).

#### Computer experiments

Learning experiments were run in Matlab using custom computer code. A first-order Euler scheme was used for the numerical integration of differential equations with an elementary time step *dt* = 0.2 ms (smaller values of *dt* had similar effects to increasing the learning rate and did not qualitatively modify our results; not shown). The membrane potentials were initially set to their resting values. Initial synaptic values were sampled from a Gaussian distribution with zero mean and standard deviation of 2. The time course of learning performance (as shown e.g. in Fig. 3) was quantified as the mean fraction *f*_*r*_ of correct decisions for relevant stimuli, and as the mean fraction *f*_*d,nr*_ of decisions taken in the presence of non-relevant stimuli. Note that perfect performance is achieved when *f*_*r*_ = 1 and *f*_*d,nr*_ = 0. Both *f*_*r*_ and *f*_*d,nr*_ were averaged across multiple training sets and then low-pass filtered according to 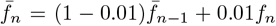, where *f*_*n*_ is the performance during stimulus exposure # *n* and 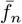 its running average up to exposure # *n*.

To ensure a fair comparison between learning curves displayed in the same figure, the initial value of the parameter *λ*^*µ*^ in Eq. 15 was selected so as to produce the same initial rate of decisions in all learning problems. This was particularly relevant when modifying the number of decision neurons (Fig. 3b-c), the number of input spike trains in each stimulus (Fig. S3), the level of input connectivity (Fig. S4), or the amount of pair-wise correlations (Fig. S5). The value of the learning rate, *η* = 10, was kept the same across all tasks.

## Acknowledgment

We acknowledge support from the National Science Foundation Grant IIS-1161852 (GLC) and the Swiss National Science Foundation through the SystemsX.ch initiative “Neurochoice” (WS). We thank Dr. Alfredo Fontanini for sharing the cortical datasets. This work is dedicated to the memory of Robert Urbanczik.

## Author contributions

GLC, RU and WS designed research. GLC supervised research. GLC, LLD and LCC performed research. GLC wrote the manuscript. All authors approved the final version of the manuscript.

## SUPPLEMENTARY INFORMATION

**Supplementary Figure 1.**
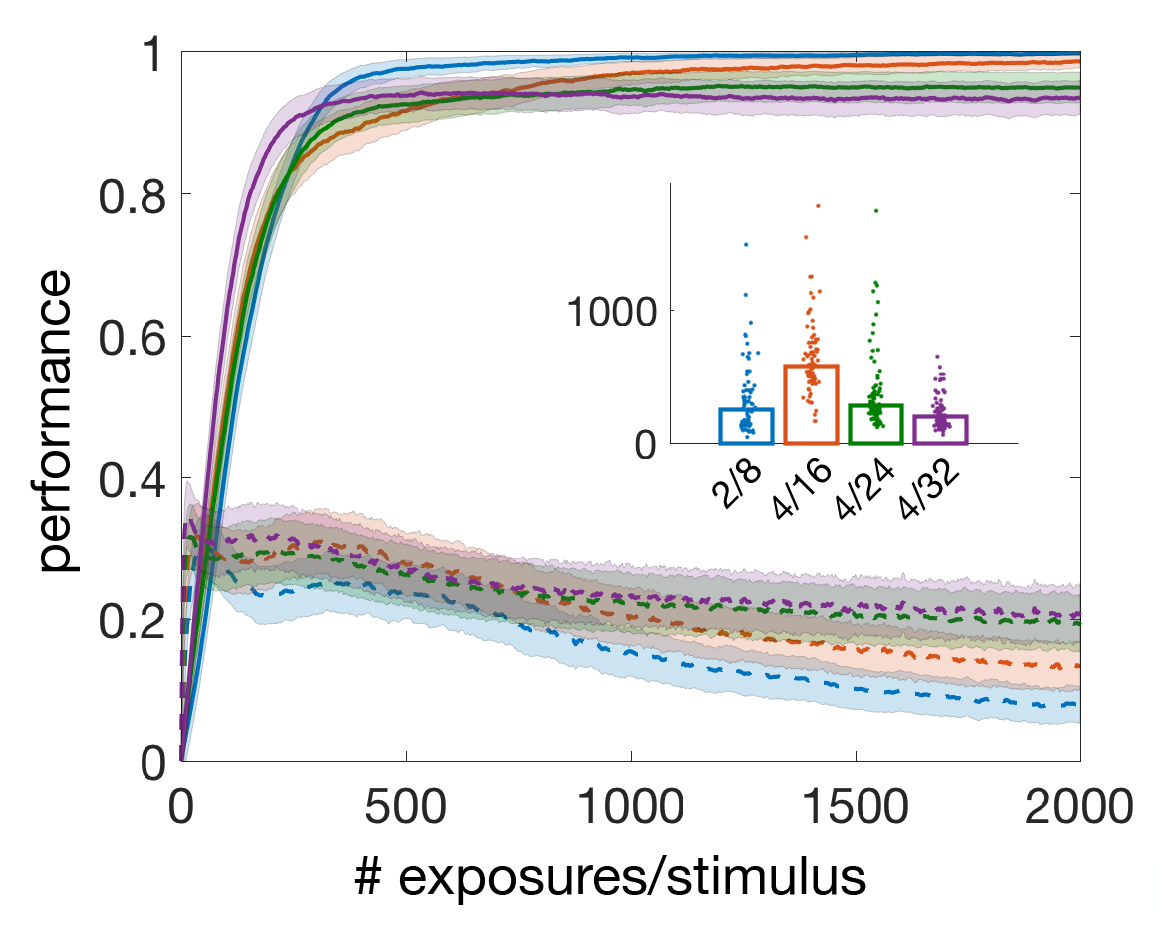
sparse stimulus embedding. Average learning curves for varying numbers of stimuli and varying ratios of non-relevant to relevant stimuli in the binary decision task (same keys as Fig. 3**b**). We trained the model with 16 (red), 24 (green) and 32 stimuli (purple). In each case, 4 stimuli were relevant (2 stimuli in each decision class), giving rise to 4:1, 6:1 and 8:1 ratios of non-relevant to relevant stimuli. The case with 8 stimuli of which 2 relevant is the same as shown in blue in Fig. 3**b** and is shown here for comparison (blue curves). Performance slightly decreased as the number of stimuli increased, but was high in all cases. We used *N* = 100 decision neurons except in the case of 32 stimuli (purple), where *N* = 200 decision neurons were used to boost performance. Mean performance with relevant stimuli raised from 85% when *N* = 100 to 94% with *N* = 200; no difference was found for non-relevant stimuli. Performance was computed in the last 1% trials of each training session and then averaged across 100 training sets for each task. Notice the faster initial learning speed with *N* = 200 due to a larger population size despite a greater number of stimuli (see Fig. 3**c**). *Inset:* Dotplots of learning thresholds for each task, with: ‘2*/*8’ = 2 relevant stimuli out of 8; ‘4*/*16’ = 4 relevant stimuli out of 16; and so on. Learning thresholds were reached in 100, 96, 88 and 83 training sets out of 100 for the task with 8, 16, 24 and 32 stimuli, respectively.

**Supplementary Figure 2.**
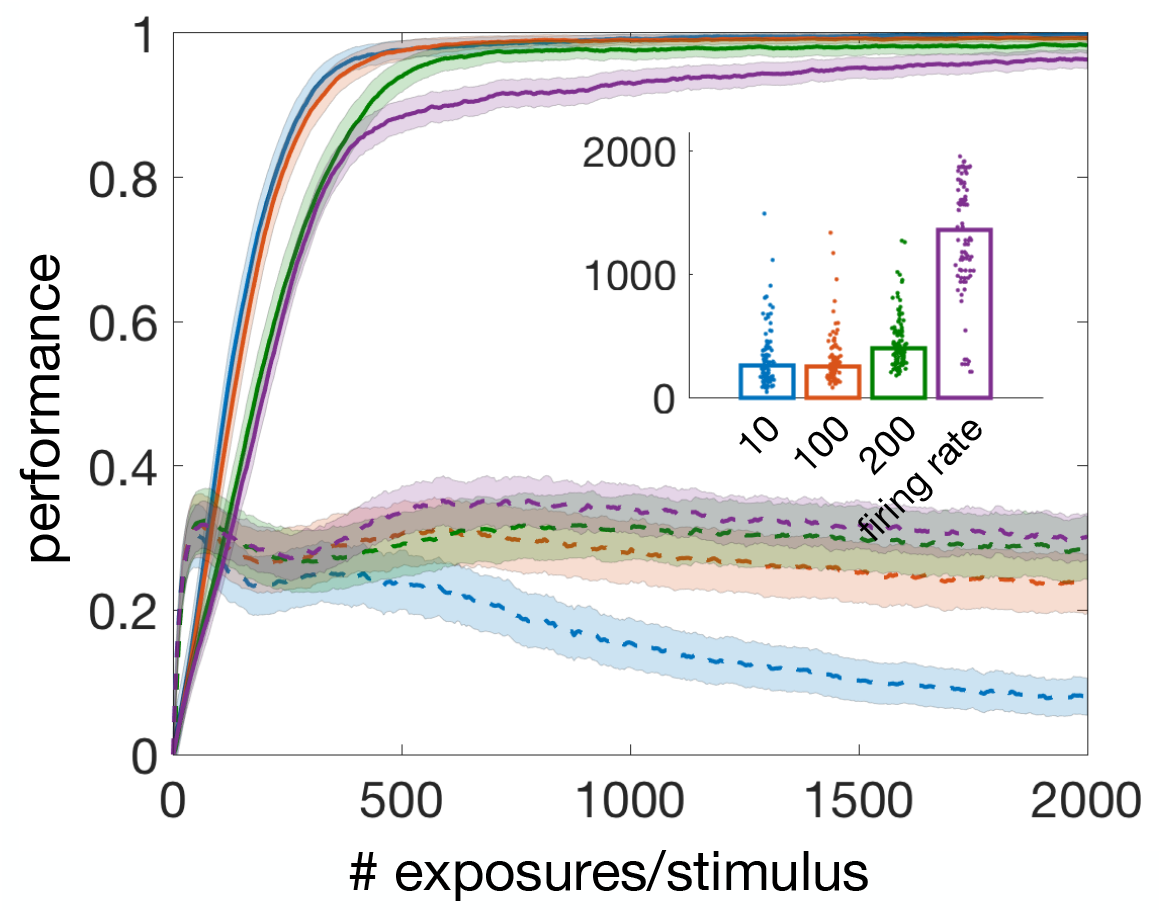
robustness to spike time jitter. Learning curves for varying jitter in the spike times of the input spike trains for the main binary decision task of Fig. 3**b** (same keys as Fig. 3**b**). Shown are the learning curves for a level of jitter *σ* = 10 ms (the main stimuli; here in blue), 100 (red), 200 (green) and firing rate coding (magenta). In firing rate coding, the spike times were generated anew at each stimulus presentation, mimicking jitter covering the entire stimulus duration. In each case we averaged across 100 learning curves except for firing rate coding where we averaged across 200 learning curves (magenta). *Inset:* Dotplots of learning thresholds for each task, with: 10, 100 and 200 being the value of *σ* in ms. Learning thresholds were always reached for *σ* ≤ 200 ms, and were reached in 65% of the training sets in the firing rate coding case. These results show that the model is tolerant to a high level of jitter in the spike times, including jitter extending over the entire duration of the stimuli. Note that 10 ms jitter and firing rate coding are, respectively, the easiest and the most difficult tasks, and therefore their learning curves (blue and magenta) bound the learning curves obtained at intermediate levels of jitter.

**Supplementary Figure 3.**
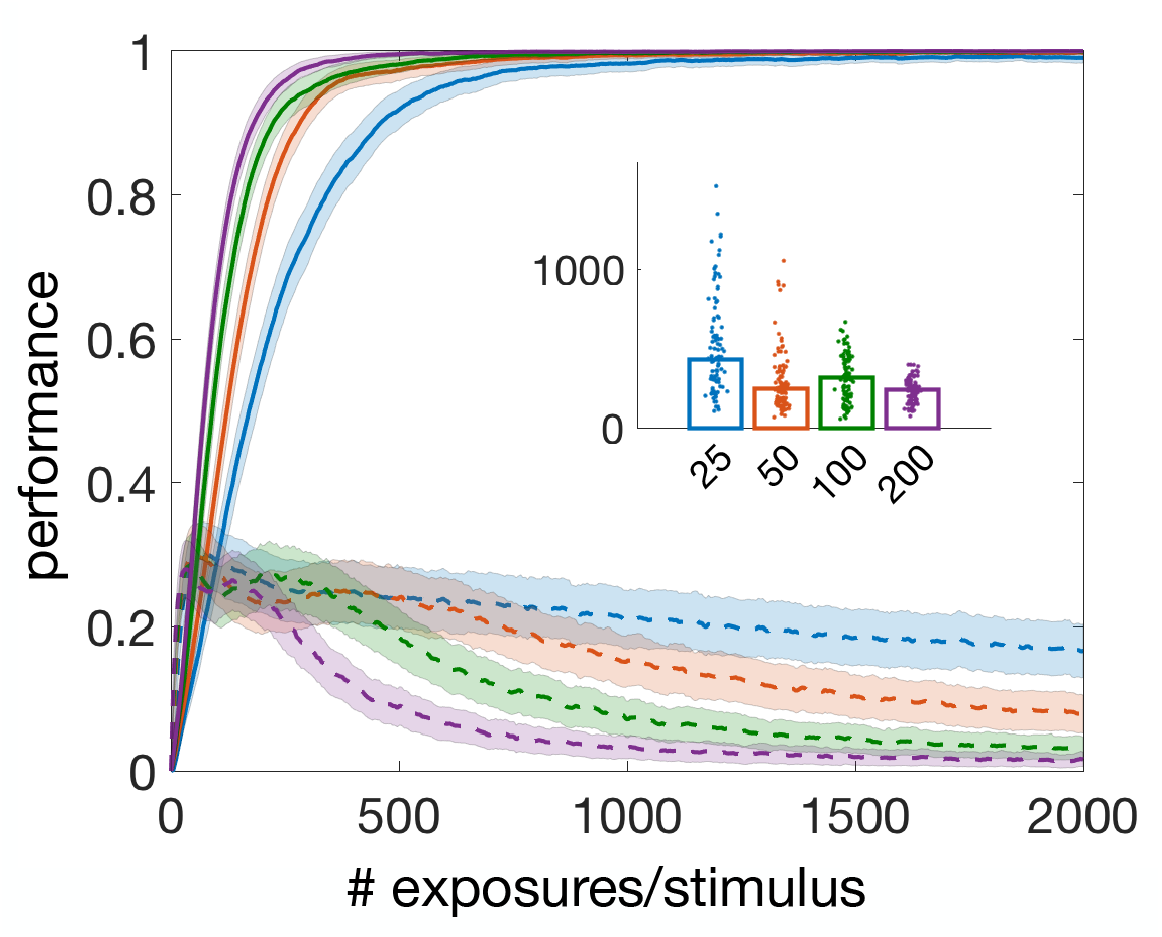
robustness to stimulus dimensionality. Learning curves for varying numbers of presynaptic spike trains for the main binary decision task of Fig. 3**b** (same keys as Fig. 3**b**). We used 25 (blue), 50 (red), 100 (green) and 200 (magenta) input spike trains and mean learning curves averaged across 100 training sets in each case. The initial value of the spike probability *λ* (see Methods, Eq. 6) was selected so as to produce an equal initial rate of decisions as the number of input spike trains varied. Final performance with relevant stimuli was high even with only 25 input spike trains, and overall learning performance improved with stimulus dimensionality. *Inset:* Dotplots of learning thresholds for each learning problem, with labels representing the number of input spike trains. Learning thresholds were reached for all training sets.

**Supplementary Figure 4.**
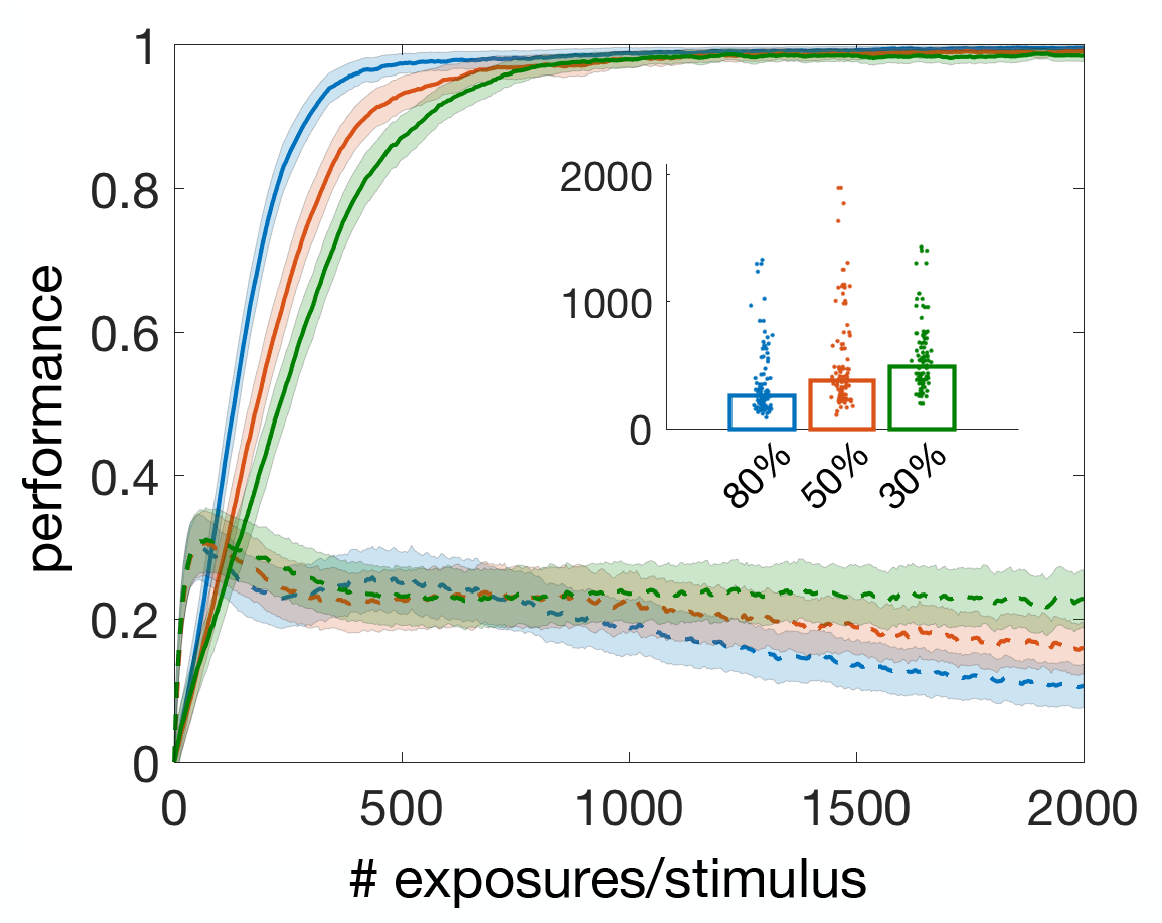
robustness to reduced connectivity. Learning curves for varying levels of input connectivity (average fraction of synaptic connections between input spike trains and decision neurons) for the main binary decision task of Fig. 3**b** (same keys as Fig. 3**b**). Connectivity values were 80% (blue), 50% (red) and 30% (green). The curve with full connectivity (blue curve in Fig. 3**b**) almost completely overlapped with the curve with 80% connectivity (not shown). To reduce the level of connectivity from 100% to *c*% *<* 100%, we removed a uniformly random fraction of (100 − *c*)*/*100 synaptic projections to the decision neurons from the 50 input spike trains. This was done independently for each prototype before training started and was then kept fixed during training in each training set. Different randomly chosen synapses were removed in different training sets. The initial value of the spike probability *λ* (see Methods, Eq. 6) was selected so as to produce an equal initial rate of decisions as connectivity varied. Decision performance after training was high also when input connectivity was sparse. *Inset:* Dotplots of learning thresholds (labels represent levels of input connectivity). Learning thresholds were reached for all training sets.

**Supplementary Figure 5.**
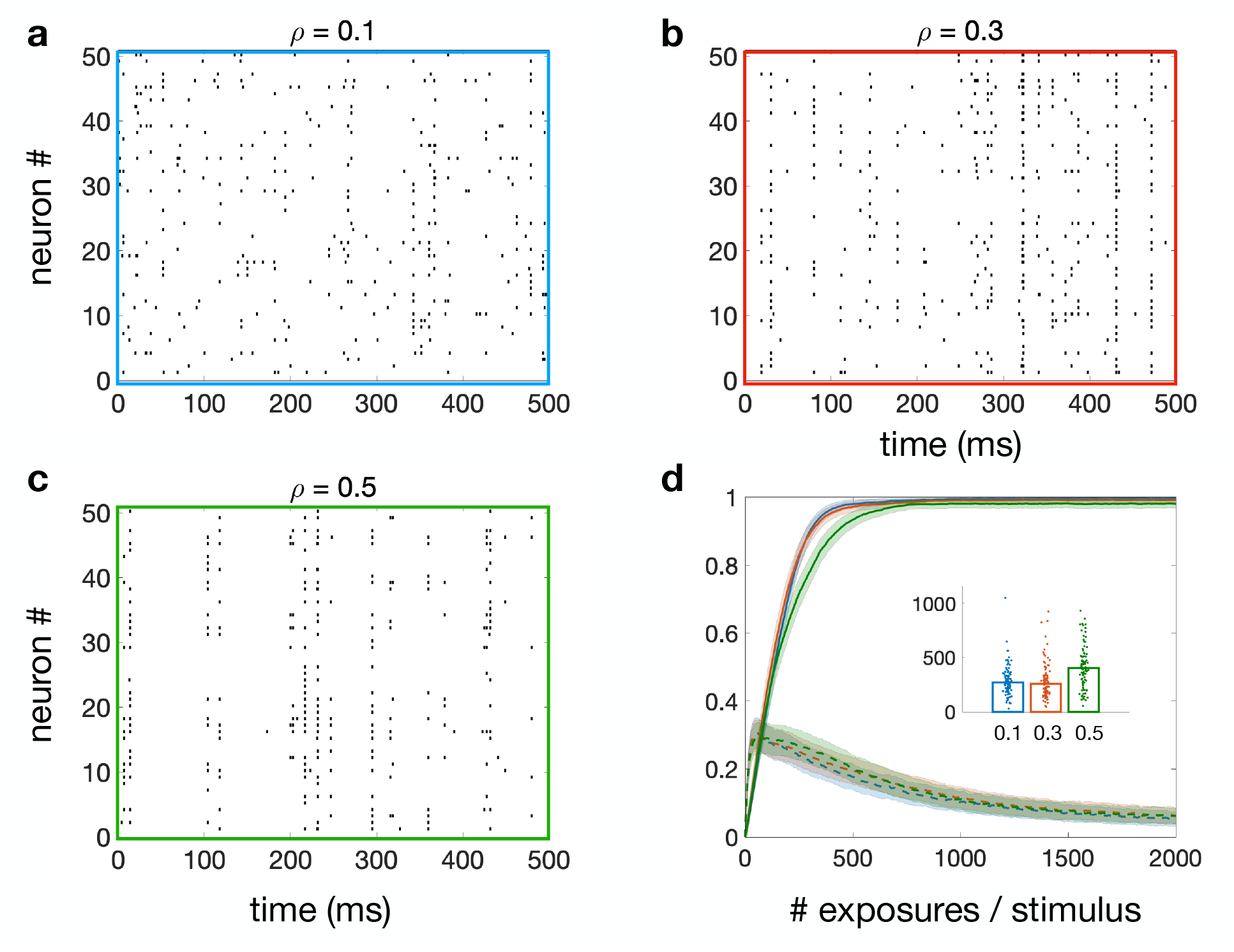
performance with correlated stimuli. Stimuli and learning curves for the main binary decision task of Fig. 3**b** with correlated input stimuli. **a, b, c)** Examples of correlated stimuli with pair-wise correlation *ρ* = 0.1, 0.3, 0.5, respectively, and firing rates uniformly distributed between 2 and 24 spikes/s (same as for the main stimuli). The spike trains were generated with the method of the dichotomized Gaussian model [18] (see Methods). **d)** Average learning curves with correlated stimuli with *ρ* = 0.1 (blue), *ρ* = 0.3 (red) and *ρ* = 0.5 (green) for the task with 2 decisions and 2 relevant stimuli out of 8 (same keys as Fig. 3b). For each value of *ρ*, 24 prototypes were generated and on each of 100 training sets, 8 out of 24 prototypes were randomly selected to be the input prototypes, jittered by 10ms on each presentation as done with the main stimuli. *Inset:* Dotplots of learning thresholds (labels represent pair-wise correlation). Learning thresholds were reached for all training sets.

**Supplementary Figure 6.**
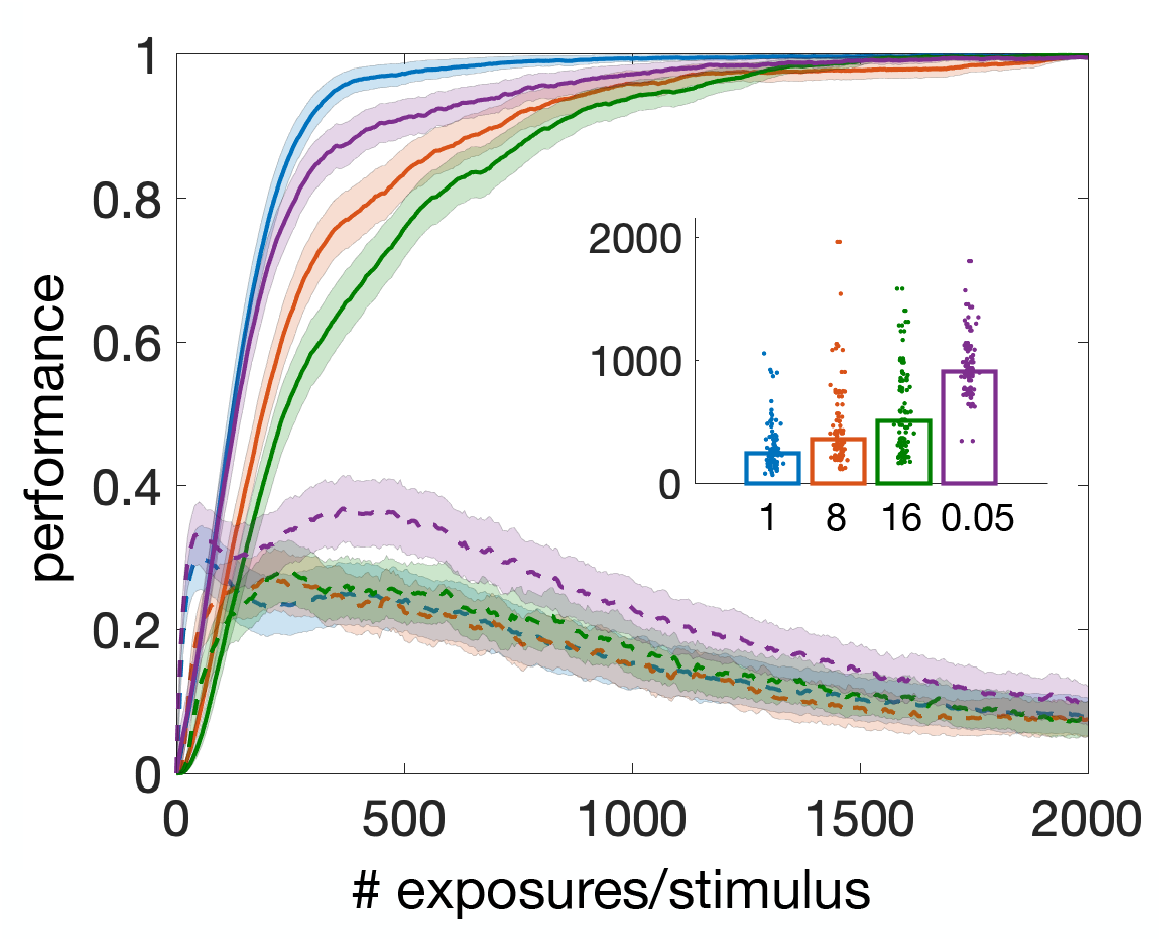
robustness of the homeostatic mechanism. Learning curves for different *mean* update times *T* of the homeostatic mechanism in the 2-way decision task of Fig. 3**b**. In each case, the update times for the spike probability (Eq. 15 of the main text) were uniformly distributed between zero and 2*T*. *T* was changed from *T* = 50 ms (magenta) to *T* = 16 s (green), with *T* = 1 s (blue) being the default value used in all other computer experiments presented in this paper. Shorter mean intervals resulted in faster learning times for *T* down to our default value of 1 s, however the performance at the end of training converged to the same level for all values of *T*. Learning with mean update times of 50 ms was slower than with *T* = 1 s, however, results could be improved by simultaneously decreasing the magnitude of change by decreasing the value of 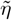 (Eq. 15 of the main text). For example, if 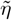 was decreased to 0.001 from 0.01, both the learning speed and the statistics of the learning thresholds with *T* = 50 ms matched that obtained with *T* = 1 s and 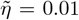 (not shown). Alternative update schedules were also tested, based on: i) intervals drawn from a Gaussian distribution, ii) constant intervals, or iii) updates occurring exactly every 32 stimulus presentations. These alternative schedules also gave similar results (not shown), confirming the robustness of the homeostatic mechanism with regard to its specific implementation and timing. *Inset:* Dotplots of learning thresholds for each experiment, with *T* (in seconds) reported under each bar. Learning thresholds were reached for all training sets.

**Supplementary Table 1.** performance and generalization with cortical datasets. Average performance for learning and generalization of the tasks with cortical data, where ‘*D*’ was the number of decisions, *P* was the number of stimuli, *P*_*rel*_ was the number of relevant stimuli, and ‘*N* ‘ was the number of decision neurons in each population. The learning curves of tasks with (*D, P*) = (2, 8), *D* = 3 and *D* = 4 are plotted in Fig. 5 of the main text. The task with *D* = 2 and *P* = 18 was used to train on 6 stimuli per class and test on one unseen stimulus per class (see the main text for details). ‘Training’: performance after training (over the last 1% portion of all stimulus presentations) across 100 training sets, reported as mean ± SD in percentage units. ‘Generalization’: generalization performance to unseen stimuli of the same class across 100 validation sets, one for each training set used for ‘training’. Each generalization run covered 10 presentations per stimulus, with learning and homeostasis disabled. ‘Control’: average performance across 100 sets on unseen stimuli of different class, used as a measure of chance generalization performance (see Methods). Performance on unseen relevant stimuli was significantly higher on stimuli of the same class (‘generalization’) compared to stimuli of different class (‘control’): p-values were *p <* 0.0004 for (*D, P*) = (2, 8), *p <* 0.015 for *D* = 3, *p <* 1.7 · 10^*−*17^ for *D* = 4, and *p <* 1.3 · 10^*−*14^ for (*D, P*) = (2, 18) (Mann-Whitney U test).

| **D** | **P** ($P_{rel}$) | **N** | **training**<br>rel. – non-rel.<br>mean(SD) – mean(SD) | **generalization**<br>rel. – non-rel.<br>mean(SD) – mean(SD) | **control**<br>rel. – non-rel.<br>mean(SD) – mean(SD) |
| --- | --- | --- | --- | --- | --- |
| 2 | 8 (2) | 100 | 99(2) – 99(1) | 48(21) – 98(3) | 37(20) – 98(6) |
| 3 | 12 (3) | 150 | 98(4) – 96(2) | 40(18) – 94(7) | 34(11) – 94(7) |
| 4 | 16 (4) | 200 | 97(5) – 93(4) | 38(13) – 84(15) | 21(10) – 83(14) |
| 2 | 18 (12) | 300 | 93(3) – 92(7) | 74(17) – 94(10) | 54(12) – 93(9) |

**Supplementary Table 2.**
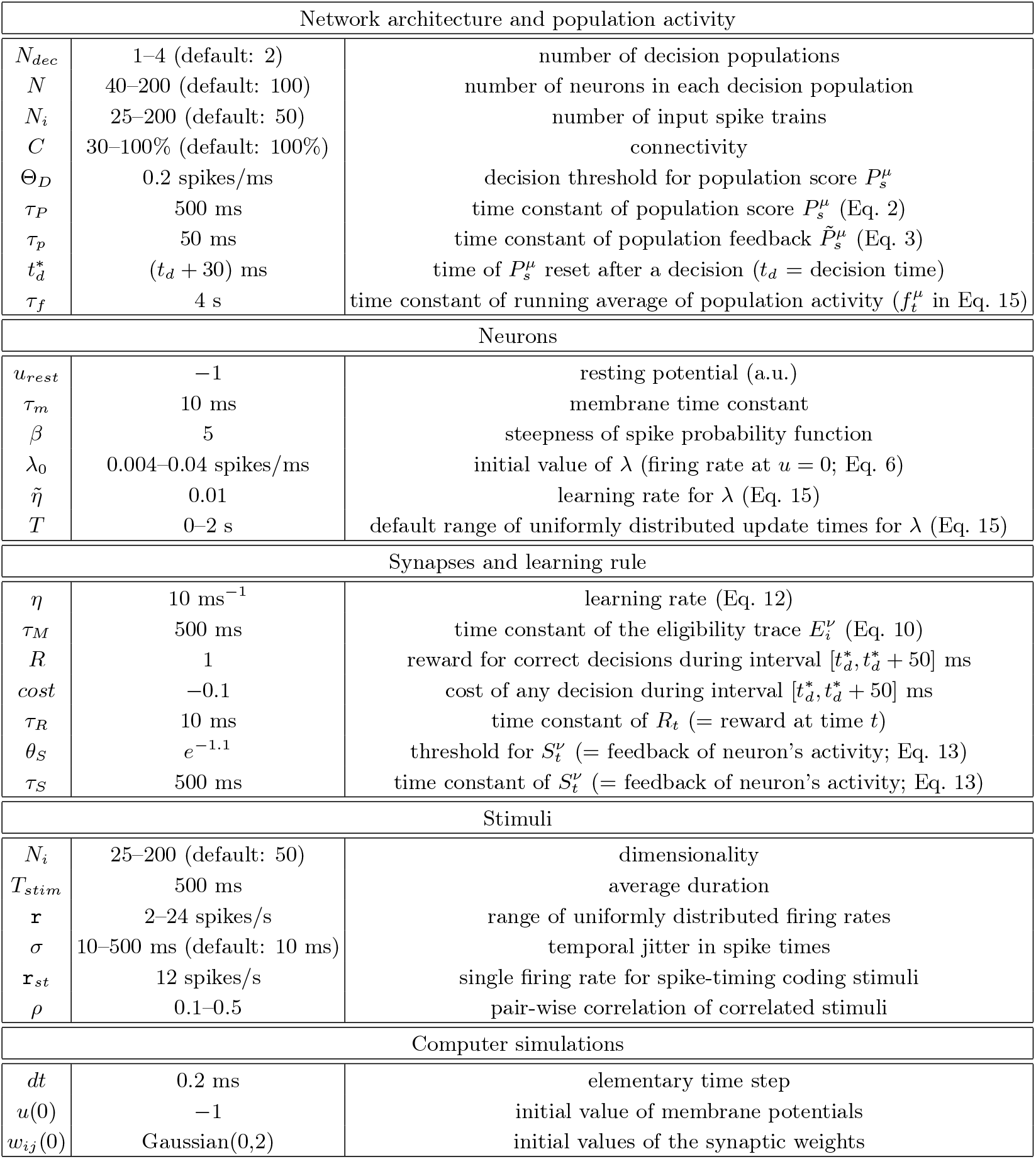
model parameters. Model and simulations parameters Values are often reported as ranges (middle column); values reported as default are the values used in most simulations.

## References

[1] P. Dayan and N. D. Daw, Cogn Affect Behav Neurosci 8, 429 (2008).

[2] X.-J. Wang, Neuron 60, 215 (2008).

[3] J. Zhang, R. Bogacz, and P. Holmes, J Math Psychol 53, 231 (2009).

[4] R. S. Sutton and A. G. Barto, Reinforcement Learning: An Introduction., 2nd ed., Adaptive computation and machine learning series (MIT Press, Cambridge, MA, 2018).

[5] D. Zeithamova, M. L. Mack, K. Braunlich, T. Davis, C. A. Seger, M. T. R. van Kesteren, and A. Wutz, J Neurosci 39, 8259 (2019).

[6] R. Urbanczik and W. Senn, Nat Neurosci 12, 250 (2009).

[7] M. Shadlen and W. Newsome, J. Neurosci. 18, 3870 (1998).

[8] M. M. Churchland, B. M. Yu, J. P. Cunningham, L. P. Sugrue, M. R. Cohen, G. S. Corrado, W. T. Newsome, A. M. Clark, P. Hosseini, B. B. Scott, D. C. Bradley, M. A. Smith, A. Kohn, J. A. Movshon, K. M. Armstrong, T. Moore, S. W. Chang, L. H. Snyder, S. G. Lisberger, N. J. Priebe, I. M. Finn, D. Ferster, S. I. Ryu, G. Santhanam, M. Sahani, and K. V. Shenoy, Nat Neurosci 13, 369 (2010).

[9] P. Dayan and L. F. Abbott, Theoretical neuroscience: computational and mathematical modeling of neural systems. (Massachusetts Institute of Technology Press, Cambridge, Mass., 2001).

[10] E. Zohary, M. N. Shadlen, and W. T. Newsome, Nature 370, 140 (1994).

[11] E. Schneidman, M. J. Berry, 2nd, R. Segev, and W. Bialek, Nature 440, 1007 (2006).

[12] F. Montani, R. A. A. Ince, R. Senatore, E. Arabzadeh, M. E. Diamond, and S. Panzeri, Philos Trans A Math Phys Eng Sci 367, 3297 (2009).

[13] I. E. Ohiorhenuan, F. Mechler, K. P. Purpura, A. M. Schmid, Q. Hu, and J. D. Victor, Nature 466, 617 (2010).

[14] S. Yu, H. Yang, H. Nakahara, G. S. Santos, D. Nikolić, and D. Plenz, J Neurosci 31, 17514 (2011).

[15] E. Ganmor, R. Segev, and E. Schneidman, Proc Natl Acad Sci U S A 108, 9679 (2011).

[16] B. Doiron, A. Litwin-Kumar, R. Rosenbaum, G. K. Ocker, and K. Josić, Nat Neurosci 19, 383 (2016).

[17] L. Mazzucato, A. Fontanini, and G. La Camera, Front. Syst. Neurosci. 10, 11 (2016).

[18] J. H. Macke, P. Berens, A. S. Ecker, A. S. Tolias, and M. Bethge, Neural Comput 21, 397 (2009).

[19] L. Mazzucato, A. Fontanini, and G. La Camera, J Neurosci 35, 8214 (2015).

[20] C. Samuelsen, M. Gardner, and A. Fontanini, Neuron 74, 410 (2012).

[21] G. Corrado and K. Doya, J. Neurosci. 27, 8178 (2007).

[22] J. I. Gold and M. N. Shadlen, Annu Rev Neurosci 30, 535 (2007).

[23] D. Lee and X.-J. Wang, in Neuroeconomics: Decision Making and the Brain, edited by P. Glimcher, C. Camerer, E. Fehr, and R. Poldrack (Academic Press, London, UK, 2009) Chap. 31, pp. 481–502.

[24] D. Green and J. Swets, Signal detection theory and psychophysics. (Wiley: New York, 1966).

[25] K. H. Britten, M. N. Shadlen, W. T. Newsome, and J. A. Movshon, J Neurosci 12, 4745 (1992).

[26] A. K. Churchland, R. Kiani, and M. N. Shadlen, Nat Neurosci 11, 693 (2008).

[27] I. M. Park, M. L. R. Meister, A. C. Huk, and J. W. Pillow, Nat Neurosci 17, 1395 (2014).

[28] A. N. Hampton, P. Bossaerts, and J. P. O’Doherty, J Neurosci 26, 8360 (2006).

[29] D. Zeithamova, W. T. Maddox, and D. M. Schnyer, J Neurosci 28, 13194 (2008).

[30] W. Schultz, J. Neurophysiol. 80, 1 (1998).

[31] A. J. Yu and P. Dayan, Neural Netw 15, 719 (2002).

[32] H. S. Seung, Neuron 40, 1063 (2003).

[33] E. M. Izhikevich, Cereb Cortex 17, 2443 (2007).

[34] N. Frémaux, H. Sprekeler, and W. Gerstner, J Neurosci 30, 13326 (2010).

[35] X. Xie and H. S. Seung, Phys Rev E Stat Nonlin Soft Matter Phys 69, 041909 (2004).

[36] J. Werfel, X. Xie, and H. S. Seung, Neural Comput 17, 2699 (2005).

[37] I. R. Fiete, M. S. Fee, and H. S. Seung, J Neurophysiol 98, 2038 (2007).

[38] J. Friedrich, R. Urbanczik, and W. Senn, Neural Comput 22, 1698 (2010).

[39] B. B. Averbeck, P. E. Latham, and A. Pouget, Nat Rev Neurosci 7, 358 (2006).

[40] M. Shadlen and W. Newsome, Current Opinion in Neurobiology 4, 569 (1994).

[41] G. Holt, W. Softky, C. Koch, and R. Douglas, J. Neurophysiol. 75, 1806 (1996).

[42] J. Friedrich, R. Urbanczik, and W. Senn, Int J Neural Syst 24, 1450002 (2014).

[43] R. Bogacz, E. Brown, J. Moehlis, P. Holmes, and J. D. Cohen, Psychol Rev 113, 700 (2006).

[44] T. Hartley, C. Lever, N. Burgess, and J. John O’Keefe, Philos Trans R Soc Lond B Biol Sci 369, 20120510 (2014).

[45] J. O’Keefe and M. L. Recce, Hippocampus 3, 317 (1993).

[46] W. E. Skaggs, B. L. McNaughton, M. A. Wilson, and C. A. Barnes, Hippocampus 6, 149 (1996).

[47] A. Ponce-Alvarez, V. Nácher, R. Luna, A. Riehle, and R. Romo, J. Neurosci. 32, 11956 (2012).

[48] L. M. Jones, A. Fontanini, B. F. Sadacca, P. Miller, and D. B. Katz, Proc Natl Acad Sci U S A 104, 18772 (2007).

[49] A. Jezzini, L. Mazzucato, G. La Camera, and A. Fontanini, J Neurosci 33, 18966 (2013).

[50] H. Shimazaki, K. Sadeghi, T. Ishikawa, Y. Ikegaya, and T. Toyoizumi, Sci Rep 5, 9821 (2015).

[51] T. Fawcett, Pattern Recognition Letters 27, 861 (2006).

[52] T. Saito and M. Rehmsmeier, PLoS One 10, e0118432 (2015).

[53] L. Rabiner, Proceedings of the IEEE 77, 257 (1989).

[54] M. Abeles, H. Bergman, I. Gat, I. Meilijson, E. Seidemann, N. Tishby, and E. Vaadia, Proc. Natl. Acad. Sci. U. S. A. 92, 8616 (1995).

[55] K. Maboudi, E. Ackermann, L. W. de Jong, B. E. Pfeiffer, D. Foster, K. Diba, and C. Kemere, Elife 7:e34467 (2018).

[56] L. Mazzucato, G. La Camera, and A. Fontanini, Nat Neurosci 22, 787 (2019).

[57] S. J. Gershman and D. M. Blei, Journal of Mathematical Psychology 56, 1 (2012).

[58] Z. Chen, Comput Intell Neurosci 2013, 251905 (2013).

[59] G. Celeux and J. Diebolt, Computational Statistics Quarterly 2, 2: 73 (1985).

[60] R. M. Neal and G. E. Hinton, in Learning in Graphical Models, edited by M. I. Jordan (Kluwer Academic Press, 1998) pp. 355–368.

[61] G. Mongillo and S. Deneve, Neural Comput 20, 1706 (2008).

[62] D. Kappel, B. Nessler, and W. Maass, PLoS Comput Biol10, e1003511 (2014).

[63] N. Frémaux, H. Sprekeler, and W. Gerstner, PLoS Comput Biol 9, e1003024 (2013).

[64] R. Legenstein, D. Pecevski, and W. Maass, PLoS Comput Biol 4, e1000180 (2008).

[65] S. Klampfl and W. Maass, J Neurosci 33, 11515 (2013).

[66] T. Masquelier, R. Guyonneau, and S. J. Thorpe, Neural Comput 21, 1259 (2009).

[67] T. Masquelier, R. Guyonneau, and S. J. Thorpe, PLoS One 3, e1377 (2008).

[68] R. Gütig, Science 351, aab4113 (2016).

[69] J. Friedrich, R. Urbanczik, and W. Senn, PLoS Comput Biol 7, e1002092 (2011).

[70] M. Rigotti, D. Ben Dayan Rubin, S. E. Morrison, C. D. Salzman, and S. Fusi, Neuroimage 52, 833 (2010).

[71] A. Genovesio, P. J. Brasted, A. R. Mitz, and S. P. Wise, Neuron 47, 307 (2005).

[72] J. Tanji, K. Shima, and H. Mushiake, Trends Cogn Sci11, 528 (2007).

[73] M. Rigotti, D. Ben Dayan Rubin, X.-J. Wang, and S. Fusi, Front Comput Neurosci 4, 24 (2010).

[74] G. La Camera, S. Bouret, and B. J. Richmond, Front Neurosci 12, 165 (2018).

[75] F. A. Mansouri, D. J. Freedman, and M. J. Buckley, Nat Rev Neurosci 21, 595 (2020).

[76] J.-P. Pfister, T. Toyoizumi, D. Barber, and W. Gerstner, Neural Comput 18, 1318 (2006).

[77] J. N. Reynolds and J. R. Wickens, Neural Netw 15, 507 (2002).

[78] M. Oram, M. Wiener, R. Lestienne, and B. Richmond, J. Neurophysiol. 81, 3021 (1999).

[79] M. C. Wiener and B. J. Richmond, Biosystems 67, 295 (2002).

[80] D. B. Katz, S. A. Simon, and M. A. Nicolelis, J Neurosci 21, 4478 (2001).

[81] N. K. Horst and M. Laubach, Front Neurosci 7, 56 (2013).

